# Mechanisms of functional compensation, delineated by eigenvector centrality mapping, across the pathophysiological continuum of Alzheimer’s disease

**DOI:** 10.1101/342246

**Authors:** Stavros Skouras, Carles Falcon, Alan Tucholka, Lorena Rami, Raquel Sanchez-Valle, Albert Lladό, Juan D. Gispert, José Luís Molinuevo

**Author notes:** **Corresponding Author** José Luis Molinuevo (MD, PhD), Barcelonabeta Brain Research Center, Pasqual Maragall Foundation. C/ Wellington 30, Barcelona E-08005, Spain.

## Abstract

**Background:** Mechanisms of functional compensation throughout the progression of Alzheimer’s disease (AD) remain largely underspecified. By investigating functional connectomics in relation to cerebrospinal fluid (CSF) biomarkers across the pathophysiological continuum of AD, we identify disease-stage-specific patterns of functional degradation and functional compensation.

**Methods:** Data from a sample of 96 participants, comprised of 49 controls, 11 preclinical AD subjects, 21 patients with mild cognitive impairment (MCI) due to AD and 15 patients with mild dementia due to AD, were analyzed. CSF ratio of phosphorylated tau protein over amyloid beta peptide 42 (p-tau/Aβ42) was computed and used as a marker of progression along the AD continuum. Whole-brain, voxel-wise eigenvector centrality mapping (ECM) was computed from resting-state fMRI and regression against p-tau/Aβ42 was performed. Surviving clusters were used as data-derived seeds in functional connectivity analyses and investigated in relation to memory performance scores (delayed free recall and memory alteration) via complementary regression models. To investigate disease-stage-specific effects, the whole-brain connectivity maps of each cluster were compared between progressive groups.

**Results:** Decreasing centrality in the inferior parietal lobule (IPL) is significantly correlated with the p-tau/Aβ42 ratio and associated to memory function impairment across the AD continuum. The thalamus, anterior cingulate (ACC), midcingulate (MCC) and posterior cingulate cortex (PCC) show the opposite effect. The MCC shows the highest increase in centrality as memory performance decays. In the asymptomatic preclinical group, MCC shows reduced functional connectivity (FC) with the left hippocampus and stronger FC with the precuneus (PCu). Additionally, IPL shows reduced FC with the cerebellum, compensated by stronger FC between cerebellum and PCC. In the MCI group, PCC shows reduced FC with PCu, compensated by stronger FC with the left pars orbitalis, insula and temporal pole, as well as by stronger FC of MCC with its anterior and ventral neighboring areas and the cerebellum. In the mild dementia group, extensive functional decoupling occurs across the entire autobiographical memory network and functional resilience ensues in posterior regions and the cerebellum.

**Conclusions:** Functional decoupling in preclinical AD occurs predominantly in AD-vulnerable regions (e.g. hippocampus, cerebellar lobule VI / Crus I, visual cortex, frontal pole) and coupling between MCC and PCu, as well as between PCC and cerebellum, emerge as intrinsic mechanisms of functional compensation. At the MCI stage, the PCu can no longer compensate for hippocampal decoupling, but the compensatory role of the MCC and PCC ensue into the stage of dementia. These findings shed light on the neural mechanisms of functional compensation across the pathophysiological continuum of AD, highlighting the compensatory roles of several key brain areas.

**Highlights:**

- Inferior parietal lobule centrality implicated in Alzheimer’s disease
- Increasing centrality in cingulate and thalamus involved in functional compensation
- Preclinical functional alterations of hippocampus compensated by precuneus
- Cerebellar involvement in functional compensation

**Abbreviations:**

Aβ42: amyloid beta peptide 42
ACC: Anterior Cingulate Cortex
AD: Alzheimer’s disease
BA: Brodmann Area
CSF: Cerebrospinal Fluid
EC: Eigenvector Centrality
ECM: Eigenvector Centrality Mapping
FC: Functional Connectivity
IPL: Inferior Parietal Lobule
MCC: Middle Cingulate Cortex
MCI: Mild Cognitive Impairment
PCC: Posterior Cingulate Cortex
PCu: Precuneus
P-tau: phosphorylated tau protein

## 1. INTRODUCTION

AD is a devastating neurodegenerative disease bolstered by a complex mechanism that has remained the subject of investigations for over a century since its initial characterization. During a tacit process of disease incubation that can last several decades, the disease remains asymptomatic. Once symptoms such as episodic memory decline, cognitive impairment and neurodegeneration become apparent, the disease is already in an advanced stage (Sperling et al., 2011). Subjects at the early stages of the pathophysiological continuum of AD maintain neuropsychological performance that is indistinguishable from healthy participants on the individual level (Dubois et al., 2016). Due to this reason, preclinical AD offers a unique window into mechanisms of functional compensation that enable the brain to exhibit resilience, reorganize adaptively and maintain unaffected functionality under the pressure of biological changes that eventually lead to neurodegeneration. fMRI methods are beginning to distinguish subtle functional changes that precede irreversible neurodegeneration (Sheline and Raichle, 2013; Leal et al., 2017; Palmqvist et al., 2017) while the brain remains able to compensate functional changes robustly (Pihlajamäki and Sperling, 2008; Teipel et al., 2015).

The cerebrospinal fluid (CSF) ratio of phosphorylated tau protein to amyloid beta peptide 42 (p-tau/Aβ42) has been used as a marker of individual pathophysiology alterations along the continuum from normal cognition to dementia due to AD (Maddalena et al., 2003). The p-tau/Aβ42 ratio has scored well as a predictor of cognitive decline in non-demented older adults (Fagan et al., 2007) and has been comparatively assessed with regards to diagnostic power across different populations (Molinuevo et al., 2013). Evidence suggested the p-tau/Aβ42 ratio as the most effective biomarker index for distinguishing dementia due to AD from semantic and frontotemporal dementias (de Souza et al., 2011). Effects of amyloid beta levels on resting state functional connectivity (FC) have recently been investigated in cognitively normal subjects (Elman et al., 2016; Palmqvist et al., 2017), suggesting effects on the intra-network and inter-network connectivity of the default mode network (DMN). In contrast to biomarkers based on measurements of only amyloid or only tau, the p-tau/Aβ42 ratio increases without saturating across the pathophysiological continuum of AD (Li et al., 2007). Thus, using the p-tau/Aβ42 ratio as a regressor of interest, enables addressing incremental and progressive connectomic changes along the entire continuum of AD, including healthy controls and the evasive preclinical phase.

Contemporary theories of network-based degradation and compensation propose that in the face of neurological insult, brain network nodes with low neural reserve degrade first while nodes with high neural reserve perform a compensatory role (Reuter-Lorenz and Cappell 2008; Seeley et al., 2009; Barulli and Stern 2013; Jacobs et al., 2013). Recent evidence shows that early β-amyloid accumulation occurs predominantly in the DMN, concurrently affecting functional connectivity in a u-shaped manner, centered around the threshold of amyloid positivity (Palmqvist et al., 2017). This is consistent with previous accounts of an initial increase of posterior DMN connectivity in the preclinical stage of AD, followed by a decrease of DMN activity in AD patients (Jack et al., 2013). Even though the DMN is traditionally associated to task-induced deactivations (Gusnard & Raichle, 2001), it is also related to attending to the external environment (Shulman et al., 1997; Raichle et al., 2001; Gilbert et al., 2007) as well as to episodic and autobiographical memory (Andrews-Hanna et al., 2010), some of the primary functions afflicted by AD. In this context, the increased functional connectivity observed in the asymptomatic stages of the pathophysiological continuum of AD, appears to reflect processes of functional compensation.

Eigenvector centrality mapping (ECM) is an assumption-free, data-driven procedure that can be performed to identify network nodes acting as hubs of connectivity and information flow (Bonacich, 1972; Borgatti, 2005; Lohmann et al., 2010). ECM can use the timeseries from all voxels to compute the eigenvector centrality (EC) of each voxel and can be particularly useful in uncovering small-world networks via functional connectivity (Koelsch & Skouras, 2014; Schoonheim et al., 2014). A previous study has shown that ECM of rs-fMRI in AD patients reveals similar patterns as FDG-PET, although less pronounced, across brain lobes (Adriaanse et al., 2016). Another pioneering study investigated changes of rs-fMRI in AD patients using voxel-wise ECM (Binnewijzend et al., 2014). In that study, EC during rs-fMRI was compared in AD patients versus their healthy family members and group differences of EC were observed in the anterior cingulate cortex (ACC) and cuneus.

Here, we investigate ECM changes with 3 Tesla fMRI in whole-brain voxel-wise resolution across the entire pathophysiological continuum of AD. By pairing the specificity of the p-tau/Aβ42 CSF biomarker with ECM, complemented by advanced normalization techniques to control for differences in brain morphology, we identify brain regions that become incrementally less central as the disease progresses and regions that concurrently compensate by becoming more central. Further testing confirms that centrality changes in these regions are implicated in memory function and the regions are used as seeds to investigate the underlying, intrinsic, progressive changes in functional connectivity (FC) that occur in each specific stage of AD. Due to its involvement in memory loss and neurodegeneration, the hippocampus and other previously established AD-vulnerable areas, e.g. inferior parietal lobule (IPL), angular gyrus, precuneus (PCu) and visual cortex (Dickerson et al., 2009) were hypothesized to be decreasing in centrality along the AD continuum. Moreover, according to the literature, we expected to observe an increase of ECM in the ACC and cuneus (Binnewijzend et al., 2014), decreased functional connectivity between the hippocampus and the PCC/PCu (Sheline and Raichle, 2013), an initial increase of posterior DMN connectivity in the preclinical stage followed by a decrease of DMN activity in patients (Jack et al., 2013) and increased connectivity in the left hippocampus, left uncus, left IPL, left thalamus, PCC and PCu, associated to previously underspecified mechanisms of cognitive compensation (Jacobs et al., 2013).

## 2. MATERIAL AND METHODS

### 2.1 Subjects

A total of 96 subjects participated in the study, at the ‘AD and other cognitive disorders unit’, Hospital Clinic i Universitari, (Barcelona, Spain): 49 controls, 11 preclinical AD subjects, 21 patients with mild cognitive impairment due to AD (MCI) and 15 patients with mild dementia due to AD. The study was approved by the local ethics committee and informed consent was obtained from all participants and caregivers in accordance with the Declaration of Helsinki. A larger number of subjects had undergone clinical and neuropsychological assessment, lumbar puncture (LP), MRI scanning, and CSF analysis. Final inclusion eligibility was determined based on: a) completeness of data (anatomical T1 data, rs-fMRI data, CSF p-tau and Aβ42 values, Apolipoprotein E genotype, neuropsychological tests and demographics); b) unambiguous diagnosis by two neurologists and one neuropsychologist; c) no comorbid clinical pathology; d) no excessive movement during functional neuroimaging.

Categorization into groups was based on the following criteria: a) Controls were CSF Aβ42 negative (over 550 pg/mL) and presented no evidence of cognitive impairment on any of the administered neuropsychological tests, in accordance with the National Institute on Aging-Alzheimer’s Association (NIA-AA) criteria (Sperling et al., 2011); b) Preclinical AD subjects presented no evidence of cognitive impairment and were CSF Aβ42 positive; c) Subjects of the MCI group were CSF Aβ42 positive, featured preserved functionality regarding daily activities, i.e. scored below 6 on the Functional Activities Questionnaire, and scored below 1.5 standard deviations from the sample mean, after accounting for the effects of age and education, on the total recall measure of the Free and Cued Selective Reminding Test (FCSRT; Grober et al., 2009) and one or more of further cognitive tests (supplementary table 1); d) Subjects of the dementia due to AD group, fulfilled the NIA-AA criteria for dementia due to AD, were CSF Aβ42 positive and in the mild stages of the disease, i.e. a score of 4 on the Global Deterioration Scale (Wesson and Luchins, 1992).

### 2.2 CSF sampling and calculation of the CSF biomarker index

All subjects underwent lumbar puncture during standard morning hours (9:00-12:00 AM). Polypropylene tubes were used to sample ten milliliters of CSF from each subject. Samples were centrifuged and stored at −80°C within one hour from collection. Using enzyme-linked immunosorbent assay kits (Furirebio-Europe, previously known as Innogenetics, Ghent, Belgium), levels of amyloid peptide Aβ42 and phosphorylated tau (p-tau) were measured. The p-tau/Aβ42 ratio, was calculated for each subject, as previously described (Maddalena et al., 2003).

### 2.3 Image acquisition

All subjects underwent MRI scanning on a 3T MRI scanner (Magnetom Trio Tim, Siemens, Erlangen, Germany). Each scanning session comprised of one high-resolution three-dimensional structural T1-weighted image acquisition (MPRAGE; TR = 2300 ms, TE = 2.98 ms, matrix size = 256 × 256, 240 sagittal slices, voxel size = 1 × 1 × 1 mm^3^) and one ten-minute resting state fMRI acquisition (300 volumes, TR = 2000 ms, TE = 16 ms, matrix size = 128 × 128, 40 axial slices, voxel size = 1.72 × 1.72 × 3 mm^3^, interslice gap = 0.75 mm). Subjects were instructed to remain still with their eyes open.

### 2.4 Image processing and analysis

To enable maximal cortical segmentation accuracy, T1 images were subjected to the N4 nonparametric nonuniform intensity normalization bias correction function (Tustison et al., 2010, 2013) of the Advanced Normalization Tools (ANTs; version 2.x, committed in January 2016; http://stnava.github.io/ANTs/; Avants et al., 2009) and to an optimized blockwise non-local means denoising filter (Coupé et al., 2008). VBM8 (Structural Brain Mapping Group, University of Jena, Jena, Germany; http://www.neuro.uni-jena.de/vbm/) and SPM12 (Wellcome Department of Imaging Neuroscience Group, London, UK; http://www.fil.ion.ucl.ac.uk/spm) were used to segment each subject’s anatomical image into grey matter, white matter and CSF. Whole-brain images with removed cranium were also derived through graph-cut (Sadananthan et al., 2010) and FSL (Analysis Group, FMRIB, Oxford, UK; https://fsl.fmrib.ox.ac.uk/fsl/) and used to compute the optimal custom anatomical template for our sample, via the ANTs multivariate template construction procedure (Avants, Tustison, et al., 2010; Avants, Yushkevich, et al., 2010) that provides optimal results in datasets with neurodegeneration (Avants et al., 2008; Avants, Yushkevich, et al., 2010). Using SyGN, neuroanatomically plausible symmetric diffeomorphic matrices were computed to transform each subject’s anatomical data to the optimal template and subsequently to MNI space (Avants, Tustison, et al., 2010; Tustison and Avants, 2013) as defined by the ICBM brain featuring high signal-to-noise ratio, sharp resolution and detailed gyrification while minimizing biases due to abnormal anatomy (Fonov et al., 2011). The cranium of the ICBM template had also been removed, similarly as for the anatomical data, prior to the normalization of datasets.

Using Matlab 2014b (MathWorks Inc. Natick, MA) and SPM12, functional data were subjected to a typical rs-fMRI preprocessing pipeline comprised of slicetime correction, estimation of movement parameters, coregistration, bandpass filtering between 0.01 and 0.1 Hz and second order detrending to remove slow signal drifts. In all subjects, movement remained below 3 mm for all translation parameters and below 1° for all rotation parameters. For further quality control, the effect of scanning-induced systematic vibration (Gallichan et al., 2010) was investigated by performing Fourier analysis on movement parameters and compensated by use of ‘scan nulling’ regressors (Lemieux et al., 2007) on affected volumes, i.e. volumes having a correlation coefficient to the mean image of the series that deviated by more than three standard deviations (r < 0.991) from the grand mean correlation coefficient for the sample (r = 0.995). Average CSF signal, average white-matter signal and 24 Volterra expansion movement parameters were regressed out of each participant’s timeseries. The five first and five last fMRI volumes of each series were discarded to avoid artefacts often produced at the edges of timeseries due to preprocessing procedures, resulting in a total of 290 functional volumes for each subject.

Each subject’s functional data were masked by the grey matter of their equivalent anatomical datasets using FSL and then normalized to MNI space in accordance with their respective diffeomorphic matrices using ANTs. Using Leipzig Image Processing and Statistical Inference Algorithms (LIPSIA; version 2.2.7 – released in May 2011; Max Planck Institute for Human Cognitive and Brain Sciences, Leipzig, Germany; http://www.cbs.mpg.de/institute/software/lipsia; Lohmann et al., 2000), each functional dataset was smoothed by a 6 mm FWHM kernel and ECM images were computed as previously described (Lohmann et al., 2010), based on positive correlations within each subject’s segmented grey matter binary mask. To enable second-level parametric inference, the resulting ECM images were gaussianized voxel-wise across all subjects (Albada and Robinson, 2007).

### 2.5 Statistical Inference

Statistical inference was performed using LIPSIA. Gaussianized ECM images from all subjects were entered into a second-level general linear model (GLM) with p-tau/Aβ42, age, gender, and education as regressors. The effect of p-tau/Aβ42 was computed, producing a z-score contrast image representing the correlations of EC with p-tau/Aβ42 in each grey matter voxel, while accounting for the effects of age, gender and education as covariates of no interest. Clusters with significant correlation between EC and p-tau/Aβ42 were identified following correction for multiple comparisons by using a combination of single-voxel probability thresholding as well as cluster-size and cluster-z-value thresholding, through 1000 iterations of Monte Carlo simulations. The initial naive cluster threshold of randomly generated maps of z-values was set to a probability level equivalent to a two-tail significance level of 0.05, because of the exploratory nature of the regression between p-tau/Aβ42 and EC. The Monte Carlo simulations accounted for anatomical priors through hemispheric symmetry and generated thresholds for cluster sizes and peak z-values given the initial threshold and the specific geometrical properties of the images (i.e. number of voxels, voxel size, spatial extent and spatial smoothness) to control for false positives and obtain the final activation maps corrected for multiple comparisons at the 0.05 level of significance (Lohmann et al., 2008).

Every ECM cluster was used as a seed cluster to calculate its standardized FC, via Fisher’s r to z transformation (Fisher 1915, 1921), with each grey matter voxel across the whole brain of each subject. Second-level GLM design matrices accounting for the effects of age, gender and education as covariates of no interest, were utilized to compute two-sample t-tests and discover voxel-wise FC differences of each ECM cluster between: a) control and preclinical groups; b) preclinical and MCI groups; c) MCI and dementia due to AD groups. Clusters featuring significant FC changes with the seed clusters were identified following correction for multiple comparisons, performed identically as for the EC clusters with the exception of setting the initial naive cluster threshold of the randomly generated maps of z-values to a probability level equivalent to a one-tail significance level of 0.05, because of the directional hypotheses regarding the FC of each expected ECM cluster in the preexisting AD literature (e.g. Sheline and Raichle, 2013; Jack et al, 2013; Jacobs et al, 2013).

To investigate whether the primary findings were associated with memory processes, we performed correlation testing between the average ECM of each seed cluster and scores on two important memory tests: delayed free recall score from the FCSRT (Grober and Buschke, 1987) and score on the memory alteration test (M@T; Rami et al., 2007). Because both memory scores showed highly significant correlations with average ECM values in most seed clusters, we repeated the main regression analysis, using identical thresholds but replacing p-tau/Aβ42 by each memory score and then computed the conjunctions with the p-tau/Aβ42 regression results, to identify the memory processes that each seed cluster was most strongly involved in.

## 3. RESULTS

### 3.1 Eigenvector centrality across the pathophysiological continuum of AD

An extensive left lateral occipital ECM cluster, encompassing parts of IPL Pgp, BA19 and BA39, correlated negatively with p-tau/Aβ42 (Table 2; Figure 1). Concurrently, three distinct clusters that correlated significantly with p-tau/Aβ42 were localized in the cingulate cortex: one in the medial cingulate cortex (MCC), primarily localized at the posterior part of BA24; one in the PCC, primarily localized at the intersection of BA31 and BA23; and one in the ACC, at the intersection between the anterior parts of BA32 and BA24 (Table 2; Figure 1). An additional cluster in the thalamus, including the ventral anterior nucleus, also correlated positively with p-tau/Aβ42 (Table 2; Figure 1).

**Figure 1.**
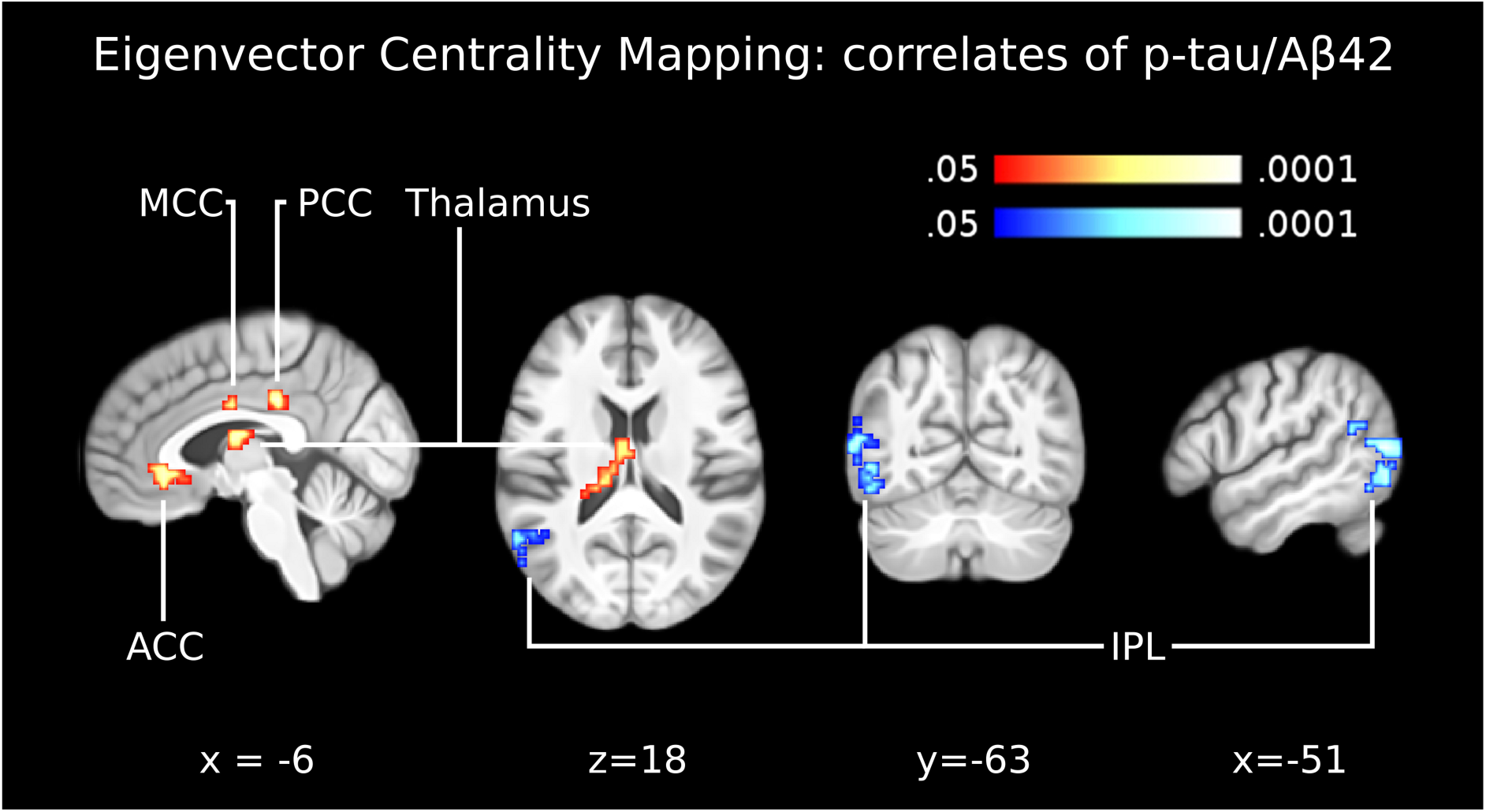
Eigenvector centrality across the pathophysiological continuum of AD. The panel shows the correlation between eigenvector centrality maps and the CSF biomarker p-tau/Aβ42, across the AD continuum from health to Alzheimer’s dementia. Clusters of centrality values significantly correlating positively with p-tau/Aβ42 were indicated in the anterior cingulate cortex (ACC), midcingulate cortex (MCC), posterior cingulate cortex (PCC) and left thalamus. One cluster of centrality values significantly correlating negatively with p-tau/Aβ42 was indicated in the inferior parietal lobule (IPL). These five clusters were used as seed regions for functional connectivity analyses (figures 2-4). Images are shown in neurological convention; all results are corrected for multiple comparisons (P < 0.05). Coordinates refer to MNI space.

**TABLE 1:** Demographics and background data. The participants’ groups have differed in terms of the relative prevalence of ApoE4 carriers. However, the differences are taken to reflect the actual prevalence of ApoE4 in Alzheimer’s disease. Due to the small number of ApoE4 carriers in our sample, including this information in the statistical models would not produce more reliable findings. Due to regional differences and historical changes in the educational system, education was modeled as an ordinal variable with a coding of “0” corresponding to no education, “1” corresponding to elementary school education, “2” corresponding to middle school education, “3” corresponding to high school education and “4” corresponding to university education.

| | Control<br>$n_a = 49$ | Preclinical AD<br>$n_b = 11$ | MCI due to AD<br>$n_c = 21$ | AD dementia<br>$n_d = 15$ | Group effect |
| --- | --- | --- | --- | --- | --- |
| **p-tau/Aβ42** | M = 0.067<br>SD = 0.02 | M = 0.158<br>SD = 0.09 | M = 0.374<br>SD = 0.17 | M = 0.382<br>SD = 0.20 | F(3,92) = 49.910<br>P < 0.001 |
| **Delayed free recall** | M = 10.729<br>SD = 1.96 | M = 8.909<br>SD = 3.11 | M = 2.5<br>SD = 3.26 | M = 2<br>SD = 2.33 | F(3,92) = 81.260<br>P < 0.001 |
| **Memory alteration** | M = 46.290<br>SD = 2.89 | M = 43.91<br>SD = 4.13 | M = 32.24<br>SD = 6.11 | M = 27.5<br>SD = 7.75 | F(3,92) = 81.179<br>P < 0.001 |
| **Age** | M = 59.816<br>SD = 6.62 | M = 68<br>SD = 7.11 | M = 69.571<br>SD = 7.92 | M = 65.667<br>SD = 9.95 | F(3,92) = 10.068<br>P < 0.001 |
| **p-tau** | M = 49.897<br>SD = 12.25 | M = 56.299<br>SD = 26.78 | M = 120.29<br>SD = 39.67 | M = 112.496<br>SD = 52.87 | F(3,92) = 35.673<br>P < 0.001 |
| **Aβ42** | M = 758.111<br>SD = 91.54 | M = 381.458<br>SD = 92.07 | M = 341.79<br>SD = 79 | M = 312.069<br>SD = 86.04 | F(3,92) = 179.707<br>P < 0.001 |
| **Education** | M = 3.082<br>SD = 0.84 | M = 2.727<br>SD = 0.79 | M = 2.476<br>SD = 1.12 | M = 2.467<br>SD = 1.25 | F(3,92) = 2.750<br>P < 0.05 |
| **Sex ratio (M / F)** | 17 / 32 | 3 / 8 | 9 / 12 | 7 / 8 | $\chi^2(3, N=96) = 14.45$<br>P = 0.694 |
| **ApoE4 carriers** | 12% | 45% | 57% | 53% | $\chi^2(3, N=96) = 14.14$<br>P < 0.001 |

**TABLE 2.** Results of the Eigenvector Centrality Mapping (ECM) correlation with p-tau/Aβ42. Entries in brackets indicate the specific areas with the highest anatomical probabilities according to the SPM Anatomy Toolbox [Eickoff et al., 2005]. The outermost right column indicates the maximal z-value of voxels within a cluster (with the mean z-value of all voxels within a cluster in parentheses). Abbreviations: MCC: Middle Cingulate Cortex; PCC: Posterior Cingulate Cortex; ACC: Anterior Cingulate Cortex; IPL: Inferior Parietal Lobule.

|  | MNI coord. | Cluster size (mm<sup>3</sup>) | z-value: max (mean) |
| --- | --- | --- | --- |
| MCC (Area 33) | 3 -6 34 | 567 | 3.31 (2.42) |
| PCC | -6 -30 37 | 540 | 3.37 (2.41) |
| L Thalamus (Temporal) | -15 -24 22 | 1080 | 3.26 (2.43) |
| ACC (Area s24/s23) | -6 30 1 | 864 | 3.01 (2.27) |
| L IPL (FG2) | -54 -72 13 | 3564 | -4.13 (-2.73) |

Average ECM in all ECM clusters correlated with FCSRT delayed free recall and M@T scores. However, the correlations survived correction for multiple comparisons only in the MCC, thalamus and IPL (Table 3). The conjunction between the regression maps for p-tau/Aβ42 and memory scores revealed a significant positive correlation of EC within the IPL cluster with FCSRT delayed free recall score and significant negative correlations of EC within the MCC and thalamus clusters with M@T score (Figure 2).

**Figure 2.**
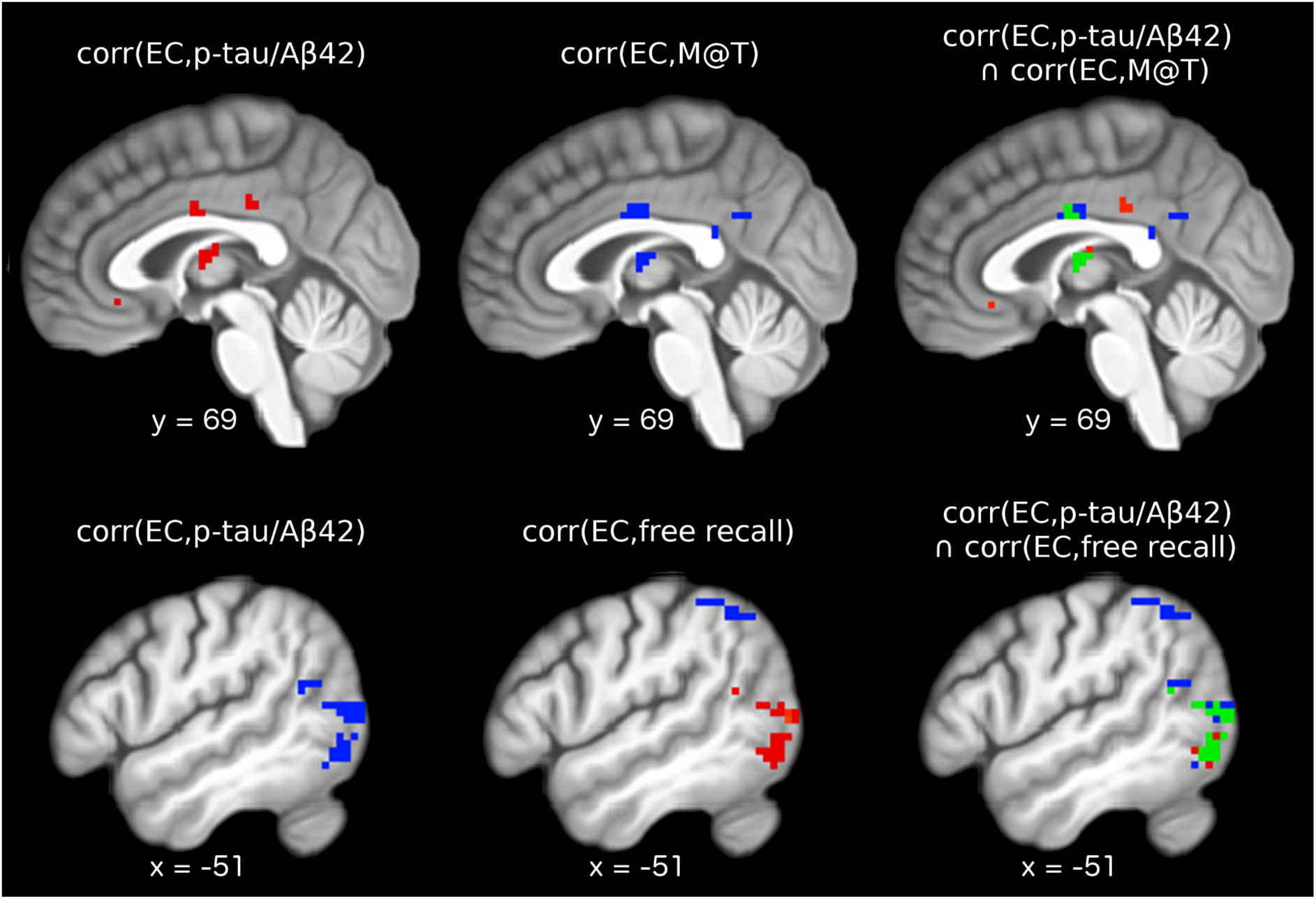
Conjunction analysis. The main eigenvector centrality regression analysis was repeated twice, replacing p-tau/Aβ42 with score on memory alteration test (M@T) and score on delayed free recall of the free and cued selective reminding test, controlling for age, gender and level of education. Results were corrected for whole-brain multiple comparisons using identical methods to the main analysis. Top row left: significant positive correlations between EC and p-tau/Aβ42; top row middle: significant negative correlations between EC and M@T scores; top row right: green voxels display regions where EC correlated significantly with both p-tau/Aβ42 and M@T scores. Bottom row left: significant negative correlation between EC and p-tau/Aβ42; bottom row middle: significant positive (red) and significant negative (blue) correlations between EC and scores of delayed free recall; bottom row right: green voxels display regions where EC correlated significantly with both p-tau/Aβ42 and delayed free recall scores.

**TABLE 3:** Correlations between average ECM values in clusters correlating with p-tau/Aβ42 and score on administered memory tests. Only correlations with p<0.001 remained significant after Bonferroni correction for multiple comparisons.

|  | MCC | PCC | Thalamus | ACC | IPL |
| --- | --- | --- | --- | --- | --- |
| Delayed free recall | r = 0.386<br>p < 0.001 | r = -0.267<br>p = 0.008 | r = -0.374<br>p < 0.001 | r = -0.232<br>p = 0.023 | r = 0.357<br>p < 0.001 |
| Memory alteration | r = -0.406<br>p < 0.001 | r = -0.2748<br>p = 0.007 | r = -0.394<br>p < 0.001 | r = -0.233<br>p = 0.022 | r = 0.392<br>p < 0.001 |

### 3.2 Functional connectivity in the preclinical AD group

In the preclinical group, compared to the control group, the ECM cluster in the IPL showed significantly lower FC with the right postcentral, precentral and supramarginal gyri (including parts of BA2, BA4 and BA3), the right frontal pole, the superior and medial frontal gyrus (primarily BA10) and an occipito-cerebellar cluster including parts of the cuneus, PCu, declive and lingual gyri bilaterally (Table 4; Figure 3, bottom row). The ECM cluster in MCC showed significantly lower FC with the left hippocampus and concurrently higher FC with the PCu in the preclinical group (Table 4; Figure 3, top row). The ECM cluster in the PCC showed significantly lower FC with the cerebellum in the preclinical group (Table 4; Figure 3, second row). The ECM clusters in the ACC and the thalamus did not show any significant differences of FC between the control and the preclinical AD groups.

**Figure 3.**
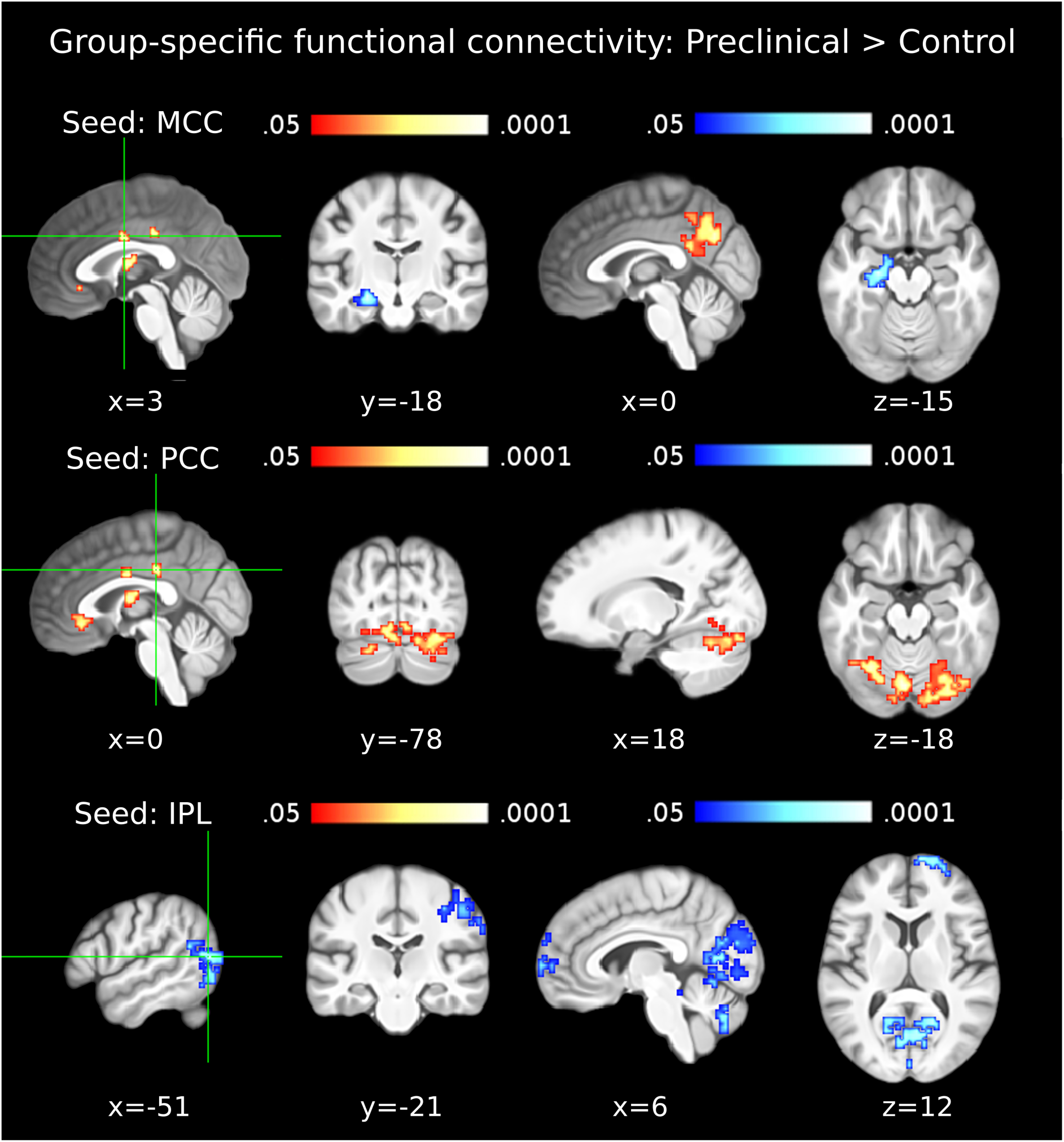
Functional connectivity in the preclinical AD group. The first column shows seed clusters derived by the data-driven ECM analysis. The results of the comparison of functional connectivity maps between preclinical and control groups (Preclinical > Control) are shown in each row, separately for three seed regions (MCC: first row, PCC: second row, IPL: third row). Each of the two remaining ECM clusters, in the ACC and thalamus, did not show functional connectivity differences between preclinical and control groups. Increases are depicted in red and decreases in blue color; images are shown in neurological convention; all results are corrected for multiple comparisons (P < 0.05); coordinates refer to MNI space.

**TABLE 4.** Results of the group-specific functional connectivity analyses for the Preclinical > Control contrast. Seed-regions used for the functional connectivity analyses were the data-driven clusters observed in the ECM results (ECM correlation with p-tau/Aβ42). For example, the MCC (which showed a positive correlation between centrality values and p-tau/Aβ42) showed decreased functional connectivity with the left hippocampus and increased functional connectivity with the precuneus in the preclinical group (compared to the control group). Entries in brackets indicate the specific areas with the highest anatomical probabilities according to the SPM Anatomy Toolbox [Eickoff et al., 2005]. The outermost right column indicates the maximal z-value of voxels within a cluster (with the mean z-value of all voxels within a cluster in parentheses). No significant differences were observed in the thalamus and ACC. Abbreviations: MCC: Middle Cingulate Cortex; PCC: Posterior Cingulate Cortex; ACC: Anterior Cingulate Cortex; IPL: Inferior Parietal Lobule; PCG: Precentral Gyrus; SMG: Supramarginal Gyrus; PCG: Postcentral Gyrus; MFG: Middle Frontal Gyrus; SFG: Superior Frontal Gyrus; SMG: Superior Medial Gyrus; HATA: Hippocampus-Amygdala Transition Area.

|  | MNI coord. | cluster size (mm<sup>3</sup>) | z-value: max (mean) |
| --- | --- | --- | --- |
| (a) MCC (Area 33) |  |  |  |
| L Precuneus; L MCC; R PCC | -6 -57 40 | 3888 | 3.81 (2.31) |
| L Hippocampus (CA1-3); HATA; Amygdala | -24 -15 -11 | 1890 | -3.93 (-2.20) |
| (b) PCC |  |  |  |
| R Cerebellar lobule VIIa crus I | 27 -69 -14 | 4806 | 3.29 (2.12) |
| L Cerebellar lobule VIIa crus | -30 -60 -14 | 2700 | 3.25 (2.08) |
| L/R Cerebellar lobule VI; Fusiform Gyrus | -6 -75 -14 | 1998 | 3.70 (2.38) |
| (c) L Thalamus (Temporal) |  |  |  |
| n.a. | n.a. | n.a. | n.a. |
| (d) ACC (Area s24/s23) |  |  |  |
| n.a. | n.a. | n.a. | n.a. |
| (e) L IPL (FG2) |  |  |  |
| R PCG (4a/4p/2/3b); SMG (PF/PFt); MFG | 48 -15 55 | 2727 | -4.31 (-2.23) |
| R/L Lingual Gyrus; Calcarine Gyrus; Cerebellum; L Cuneus | -15 -66 19 | 25569 | -3.90 (-2.26) |
| R SFG (Fp1); SMG (Fp1/Fp2) | 9 69 16 | 2484 | -3.64 (-2.26) |

### 3.3 Functional connectivity in the MCI due AD group

In the MCI due to AD group, compared to the preclinical AD group, the ECM cluster in the PCC showed significantly lower FC with PCu and concurrently higher FC with the right insula (Table 5; Figure 4, second row). Additionally, the MCC cluster showed significantly higher FC with the middle part of BA24 and cerebellum in the MCI group (Table 5; Figure 4, top row). The ECM clusters in the IPL, ACC and the thalamus did not show any significant differences of FC between the preclinical and MCI groups.

**Figure 4.**
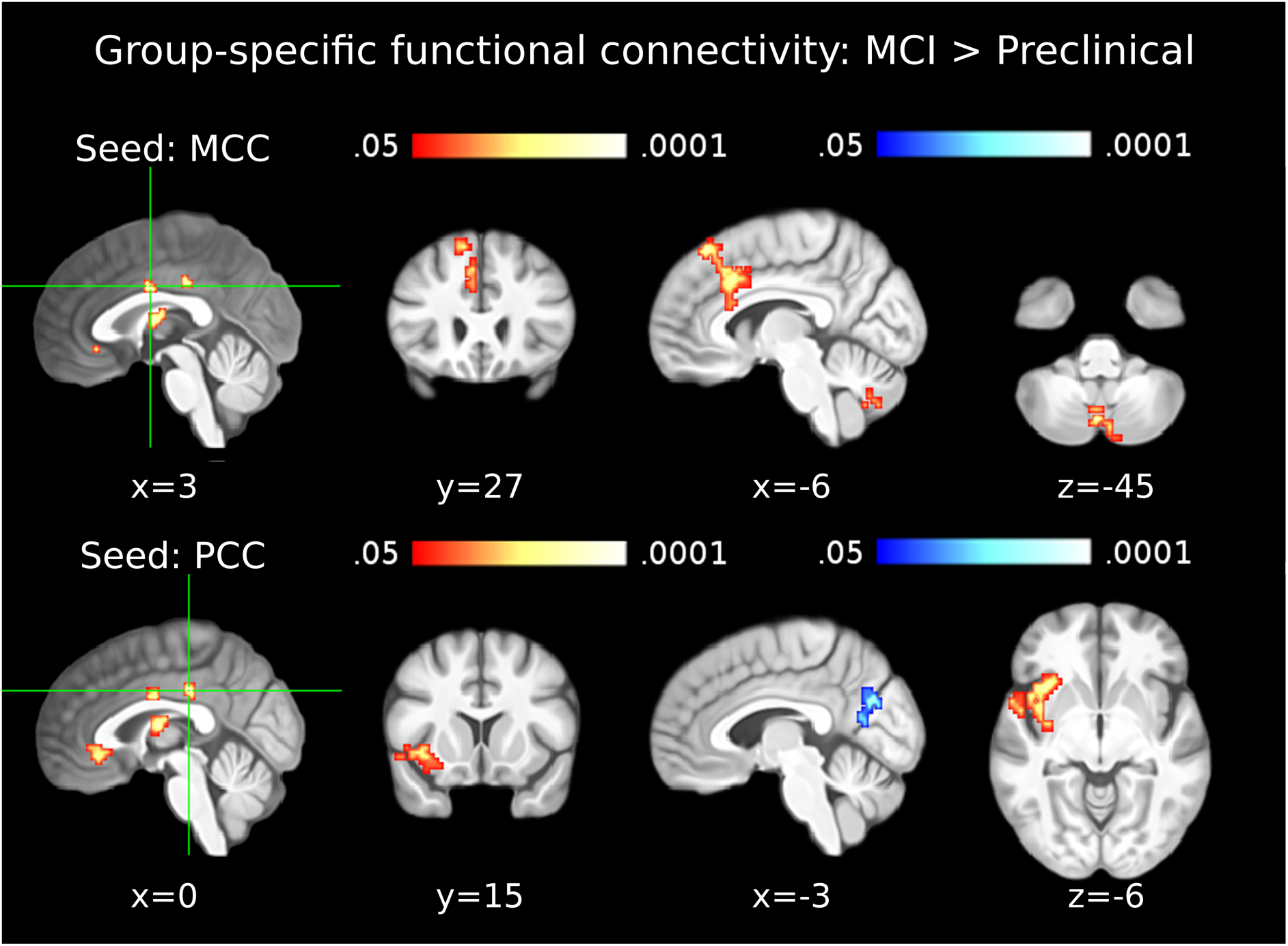
Functional connectivity in the MCI due to AD group. The first column shows seed clusters derived by the data-driven ECM analysis. The results of the comparison of functional connectivity maps between mild cognitive impairment (MCI) and preclinical groups (MCI > Preclinical) are shown in each row, separately for two seed regions (MCC: first row, PCC: second row). Each of the three remaining ECM clusters, in the ACC, thalamus and IPL, did not show functional connectivity differences between MCI and preclinical groups. Increases are depicted in red and decreases in blue color; images are shown in neurological convention; all results are corrected for multiple comparisons (P < 0.05); coordinates refer to MNI space.

**TABLE 5.**
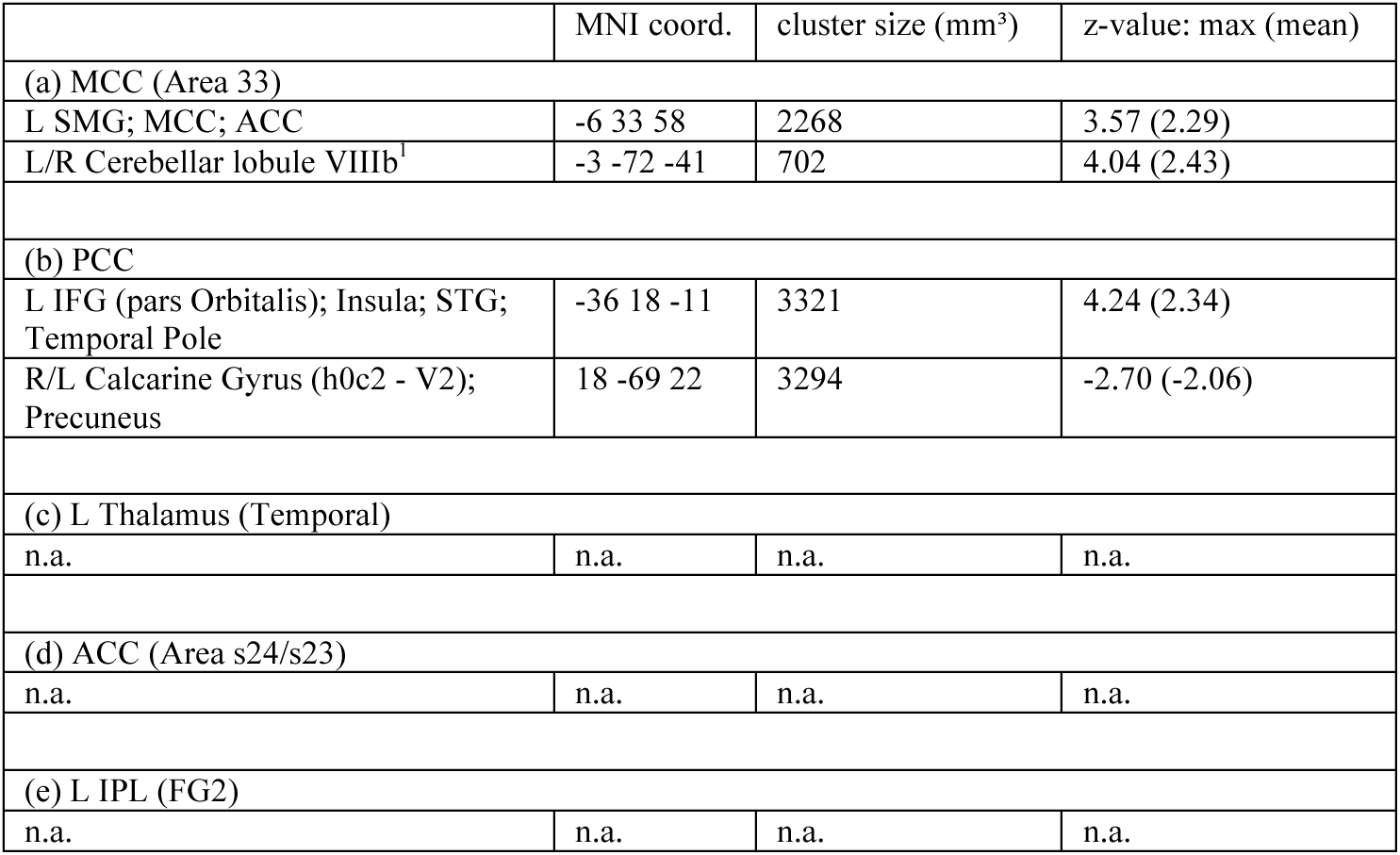
Results of the group-specific functional connectivity analyses for the MCI > Preclinical contrast. Seed-regions used for the functional connectivity analyses were the data-driven clusters observed in the ECM results (ECM correlation with p-tau/Aβ42). Entries in brackets indicate the specific areas with the highest anatomical probabilities according to the SPM Anatomy Toolbox [Eickoff et al., 2005]. The outermost right column indicates the maximal z-value of voxels within a cluster (with the mean z-value of all voxels within a cluster in parentheses). No significant differences were observed in the thalamus, the ACC and the IPL. Abbreviations: MCC: Middle Cingulate Cortex; PCC: Posterior Cingulate Cortex; ACC: Anterior Cingulate Cortex; IPL: Inferior Parietal Lobule; SMG: Supramarginal Gyrus; IFG: Inferior Frontal Gyrus; STG: Superior Temporal Gyrus.

|  | MNI coord. | cluster size (mm<sup>3</sup>) | z-value: max (mean) |
| --- | --- | --- | --- |
| (a) MCC (Area 33) |  |  |  |
| L SMG; MCC; ACC | -6 33 58 | 2268 | 3.57 (2.29) |
| L/R Cerebellar lobule VIIIb<sup>1</sup> | -3 -72 -41 | 702 | 4.04 (2.43) |
| (b) PCC |  |  |  |
| L IFG (pars Orbitalis); Insula; STG; Temporal Pole | -36 18 -11 | 3321 | 4.24 (2.34) |
| R/L Calcarine Gyrus (h0c2 - V2); Precuneus | 18 -69 22 | 3294 | -2.70 (-2.06) |
| (c) L Thalamus (Temporal) |  |  |  |
| n.a. | n.a. | n.a. | n.a. |
| (d) ACC (Area s24/s23) |  |  |  |
| n.a. | n.a. | n.a. | n.a. |
| (e) L IPL (FG2) |  |  |  |
| n.a. | n.a. | n.a. | n.a. |

### 3.4 Functional connectivity in the dementia due to AD group

In the dementia due to AD group, compared to the MCI due to AD group, the IPL cluster showed significantly lower FC with the middle and superior frontal gyri, including ACC, BA9, BA32, BA10, BA11, as well as the right middle and superior temporal gyri, BA21 and the superior temporal pole (Table 6; Figure 5, bottom row). Concurrently, significantly higher FC was observed between the IPL the cerebellum (declive, culmen and vermis lobules VI-VIII).

**Figure 5.**
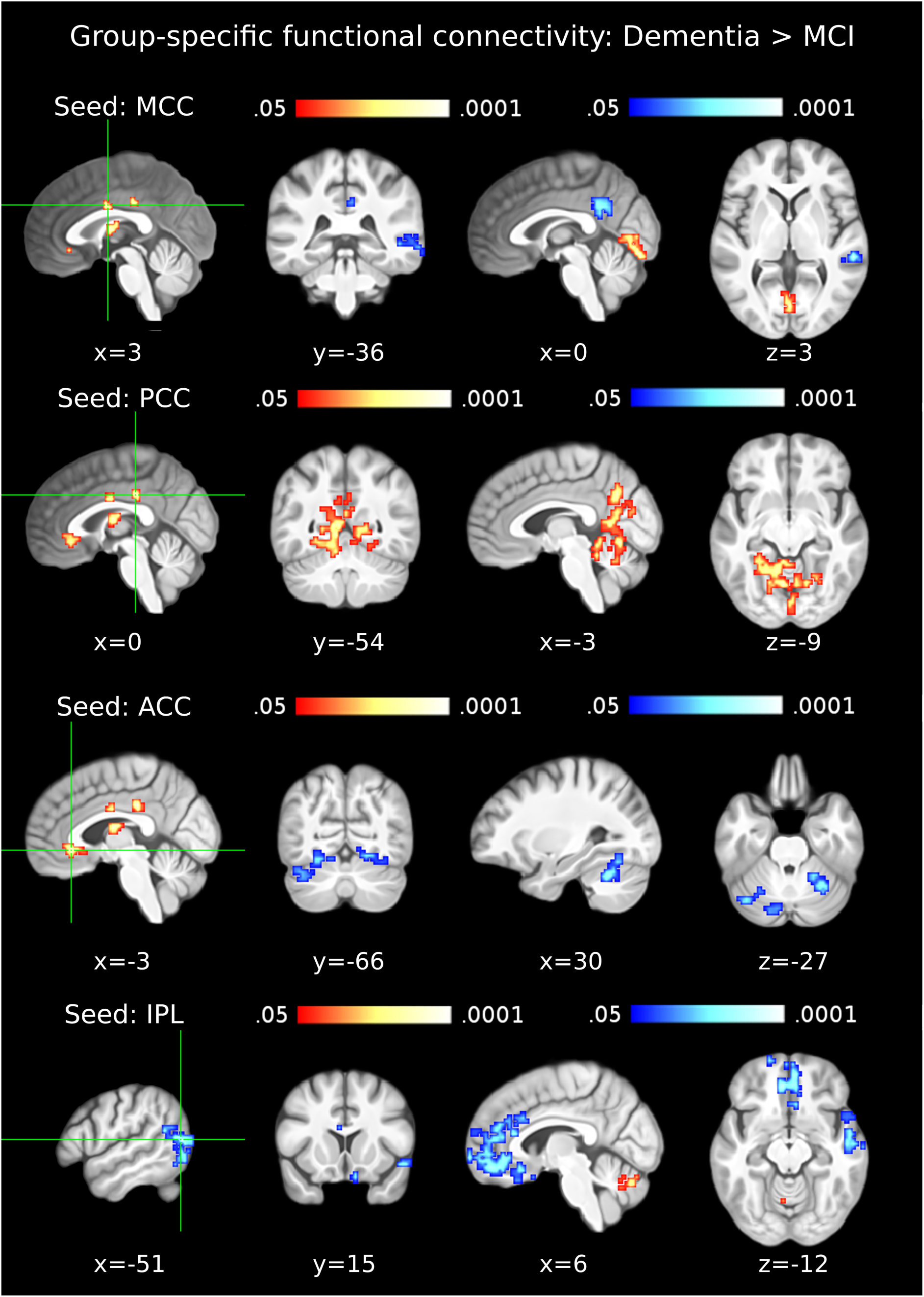
Functional connectivity in the dementia due to AD group. The first column shows seed clusters derived by the data-driven ECM analysis. The results of the comparison of functional connectivity maps between dementia and MCI due to AD (Dementia > MCI) are shown in each row, separately for four seed regions (MCC: first row, PCC: second row, ACC: third row, IPL: fourth row). The remaining ECM cluster, in the thalamus, did not show functional connectivity differences between dementia and MCI groups. Increases are depicted in red and decreases in blue color; images are shown in neurological convention; all results are corrected for multiple comparisons (P < 0.05); coordinates refer to MNI space.

**TABLE 6.**
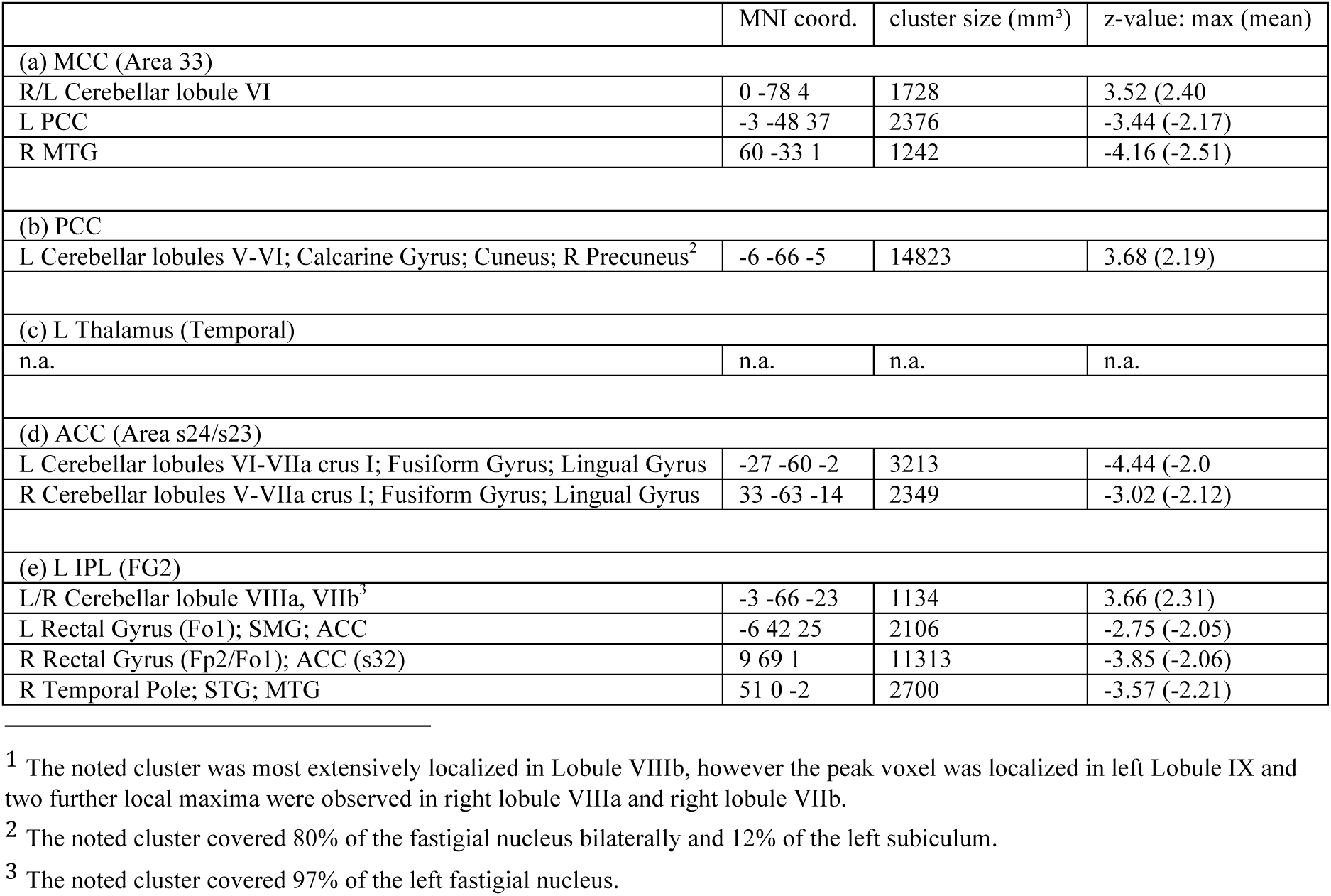
Results of the group-specific functional connectivity analyses for the Dementia > MCI contrast. Seed-regions used for the functional connectivity analyses were the data-driven clusters observed in the ECM results (ECM correlation with p-tau/Aβ42). Entries in brackets indicate the specific areas with the highest anatomical probabilities according to the SPM Ana tomy Toolbox [Eickoff et al., 2005]. The outermost right column indicates the maximal z-value of voxels within a cluster (with the mean z-value of all voxels within a cluster in parentheses). No significant differences were observed in the thalamus. Abbreviations: MCC: Middle Cingulate Cortex; PCC: Posterior Cingulate Cortex; ACC: Anterior Cingulate Cortex; IPL: Inferior Parietal Lobule; MTG: Middle Temporal Gyrus; SMG: Supramarginal Gyrus; STG: Superior Temporal Gyrus; SMG: Superior Medial Gyrus.

The MCC cluster showed significantly lower FC with the PCu, posterior cingulate cortex (PCC) and right temporal gyrus in the dementia due to AD group and concurrently increasing FC with the lingual gyri bilaterally (Table 6; Figure 5, top row). The ACC showed significantly lower FC with the cerebellum, the fusiform gyri bilaterally and the lingual gyri bilaterally in the dementia due to AD group compared to the MCI due to AD group (Table 6; Figure 5, third row). The PCC cluster showed significantly higher functional connectivity with the PCu, cerebellum and lingual gyri bilaterally in the dementia due to AD group (Table 6; Figure 5, second row). The ECM cluster in the thalamus did not show any significant differences of FC between the MCI and dementia groups.

## 4. DISCUSSION

### 4.1 Eigenvector centrality across the pathophysiological continuum of AD

The extensive left lateral occipital cluster encompassing IPL Pgp, BA19 and angular gyrus (BA39) that correlated negatively with p-tau/Aβ42, is known to be an area vulnerable to regional cortical thinning in AD (Dickerson et al., 2009; Fortea et al., 2011) and a meta-analysis suggests the BA39-BA19 intersection to be important in episodic auto-biographical memory (Martinelli et al., 2013). In healthy young adults, the identified area features the maximal degree of connectivity with the rest of the brain and is considered to be the foremost cortical hub with the highest metabolic demands (Buckner et al., 2009). Beyond the observed correlation of EC in the IPL with FCSRT delayed free recall score, the IPL is serving an important role in further memory processes, as well as in the flexible coupling between the default mode network and the frontoparietal control network subserving autobiographical planning (Spreng et al., 2010). Therefore, increasing EC in the cingulate cortex and thalamus presumably represents a compensatory response to the functional degradation of IPL and memory performance. The following evidence corroborates this interpretation.

Meta-analyses show the particular parts of MCC (posterior BA24) and ACC (BA32/BA22 junction) that correlate positively with p-tau/Aβ42, to be involved in working memory (Wager and Smith, 2003), as well as the PCC cluster (BA31; BA23) to be involved in self-autobiographical memory (Martinelli et al., 2013). The significant correlation between p-tau/Aβ42 and EC in the thalamus points to the crucial potential of strategic infarcts of the thalamus, e.g. due to vascular causes, to perform an instantaneous causal role in the manifestation of dementia (Szirmai et al., 2002). A recent review emphasized the importance of the anterior nucleus of the thalamus in the manifestation of AD (Aggleton et al., 2016) and although the thalamic cluster did not show significant FC differences between consecutive phases of AD, previous evidence suggest decreased FC of thalamus with the DMN and increased FC of thalamus with the temporal lobe, prefrontal cortex (BA11) and the BA5-BA7 junction in AD patients (Zhou et al., 2013). An additional meta-analysis illustrated a consistent pattern of robust modulations between the DMN and task-related activation in the thalamus (Jacobs et al., 2013), suggesting that FC of the thalamus may differ between groups mostly during task-related activity. During task-free fMRI, it is likely that the changes of FC between each consecutive stage of AD are not as drastic and therefore remain below statistical significance.

### 4.2 Functional connectivity in the preclinical AD group

FC differences of hippocampus and BA24 had previously been observed only between control and MCI groups (Wang et al., 2006). Our results suggest that the onset of the functional decoupling between the hippocampus and BA24 occurs earlier than previously believed, during the preclinical stage. Given no significant alterations in cognitive performance during the preclinical stage (Sperling et al., 2011), the concurrent functional coupling of the exact same BA24 cluster with the PCu emerges as a compensatory mechanism. The PCu, possesses extensive connectivity with several other important brain areas (Cavanna and Trimble, 2006), including the hippocampus, enabling it to compensate hippocampal decoupling from BA24. The meta-analysis of fMRI studies in AD is also aligned with the supposition that the PCu plays a compensatory role during the early stages of the disease, prior to any perceivable cognitive decline (Jacobs et al., 2013). The functional purpose of the increased FC between PCu and BA24, given a functional decoupling between BA24 and hippocampus, could be related to maintaining spatial navigation abilities (Cavanna and Trimble, 2006; Ghaem et al., 1997) and episodic autobiographical memory (Martinelli et al., 2013; Söderlund et al., 2012). With regards to the latter, it is also plausible that the concerned FC increase is further compensating the preclinical decoupling between PCu and IPL (Figure 3; Table 4).

During the preclinical stage, the IPL decouples not only from the PCu but also from the cuneus, cerebellum, lingual gyri, fusiform gyri, the frontal pole and a cluster containing parts of the right precentral, postcentral and supramarginal gyri. The FC decrease of the IPL with the cerebellum, lingual gyri and fusiform gyri is compensated by FC increases of PCC with the cerebellum, linguaI gyri and fusiform gyri (Figure 3; Table 4). However, the FC decrease of IPL with the part of the occipital cluster localized in the primary and secondary visual cortices remains uncompensated. Our findings suggest that the FC decrease of the IPL with the visual cortex is prolonged through the subsequent stages of AD, plausibly relating to AD-characteristic regional cortical thinning of both the IPL and the visual cortex (Dickerson et al., 2009). Further areas with early uncompensated FC decreases are the right lateral cluster spreading between the postcentral, precentral and supramarginal gyri and the cluster located in the frontal pole. Collectively, the uncompensated decreases of FC in the preclinical stage correspond to the neural correlates of subtle cognitive effects accounting for changes in global cognitive ability (Bäckman et al., 2005).

To our knowledge, this is the first evidence of cerebellar FC changes occurring in the preclinical stage of AD. Similar cerebellar subregions to the ones identified in our study, specifically Crus I/II and lobule VI, are targeted by atrophy in AD and frontotemporal dementia, respectively (Guo et al., 2016). Although the exact details of these patterns remain to be clarified, current evidence suggest FC of Crus I/II predominantly with the executive network and of lobule VI with the salience network (Habas et al., 2009; Skouras et al., 2018). Additionally, both cerebellar regions have been found to correlate with the anterior prefrontal cortex (Krienen and Buckner, 2009). In the present study, noticeable overlap between the clusters shown in figure 3 (second row and third row) was observed in lobule VI/Crus I (supplementary figure S1).

### 4.3 Functional connectivity in the MCI due to AD group

As AD progresses into the MCI stage, characterized by hippocampal neurodegeneration (Gispert et al, 2015), the PCu can no longer capacitate compensatory support for hippocampal decoupling. This is evident from the functional decoupling of PCu/cuneus from PCC (Table 5) corroborating previous evidence of lower FC in the PCu during the MCI stage (Binnewijzend and Schoonheim, 2012) and supporting models of preclinical compensation and clinical degradation in the PCu (Jack et al., 2013; Jacobs et al., 2013). In response, functional coupling of PCC with the insula, BA47, superior temporal gyrus and left temporal pole emerges as a tertiary compensatory mechanism (Figure 4; Table 5). Both the insula and BA47 are involved in episodic and working memory (Kapur et al., 1994; Kurth et al., 2010; Ranganath et al., 2003). Moreover, the left insula is part of a network related to spatial navigation (Cavanna and Trimble, 2006; Ghaem et al., 1997) and the specific left inferio-anterior region of the insula observed in the present study, is also involved in processing interoception, emotions and empathy (Kurth et al., 2010). It is plausible that the insula is utilizing its capacity to rapidly switch between internal mentation and external attention (Menon and Uddin 2010) to compensate for the functionality of the frontoparietal control network (Spreng et al., 2010) that is affected more drastically by the ensuing recession of IPL centrality.

During the MCI stage of AD, functional coupling of MCC with the cerebellum, the posterior part of BA32 and the left BA8 can account for the emergence of new cognitive strategies that MCI subjects employ in order to compensate memory and cognitive impairments and manage to cope with their daily activities. In particular, the observed subregion of the cerebellum supports executive function (Habas et al., 2009) and has been observed to interact with the hippocampus during sequence-based navigation (Iglόi et al., 2014). The observed subregion of BA32 subserves working memory (Hill et al., 2014; Wager and Smith, 2003; Zhang et al., 2003), while BA8 appears to be involved in working memory (Babiloni et al., 2005) and processing related to uncertainty (Volz et al., 2004). Additional evidence suggests that BA8 is one of the main loci of autobiographical memory (Janata, 2009) and significantly more so in borderline personality disorder patients (Schnell et al., 2007), a population that also suffers from reduced hippocampal volume (Driessen et al., 2000) similarly to the MCI group due to AD (Gispert et al., 2015).

### 4.4 Functional connectivity in the dementia due to AD group

As expected, extensive and uncompensated functional decoupling is observed in the stage of dementia due to AD (Figure 5; Table 6). The right BA21, involved in prosody processing (Ethofer et al., 2006; Hesling et al., 2005) and more importantly in memory integration (Backus et al., 2016), decouples from both the IPL and MCC. The IPL and MCC additionally decouple from the ACC and PCC respectively, exhibiting in this stage for the first time an anti-correlation pattern among ECM clusters. Even though most ECM clusters remain central hubs in dementia due to AD, they seem less coherent as a network than in earlier stages of the disease. Moreover, the lingual gyrus decouples from ACC but increases FC with MCC and PCC, continuing the compensatory effort commenced during the preclinical stage.

Certain portion of the PCu also appears to remain relatively adaptive during dementia due to AD. Although the increased FC of the PCu with the MCC that was observed in the preclinical stage decays during the final stage of AD, a more posterior, marginally overlapping, portion of the PCu still increases FC with the PCC (see supplementary figure S2). Meta-analysis showed that alterations due to AD in the PCu are functionally heterogeneous, with anterior and posterior parts associated to self-referential thoughts and episodic memory retrieval respectively, as well as contributing differentially to compensatory and neurodegenerative processes (Jacobs et al., 2013). The increased FC of the MCC with BA18 can be responsible for the previously observed dis-inhibition of saccades suggested by a recent review (Molitor et al., 2015) due to the involvement of BA18 in the saccade network (McDowell et al., 2008).

The functional decoupling of the middle temporal gyrus, superior temporal gyrus and BA21 from both the IPL and the MCC (BA24), in combination with the decoupling of the IPL from an extensive medial frontal cluster that includes ACC, BA32, BA9, BA10 and BA11, remain uncompensated and appear to be underpinning the dysfunctional state of dementia due to AD. Indeed, BA21 and BA22 are involved in the crucial function of memory integration (Backus et al., 2016). Moreover, the observed frontal cluster engulfs the neural substrates of the self-memory system subserving episodic autobiographical memory, semantic autobiographical memory and memory of the conceptual self (Martinelli et al., 2013). The extensive functional decoupling of these areas with the IPL is illustrating the final collapsing of memory functions in dementia due to AD.

Overall, in dementia due to AD, the cerebellum, the PCC, the PCu and the lingual gyri continue to show increases in connectivity. In this regard and in addition to the ECM clusters, these brain areas appear to be spared by AD and emerge as local maxima of neural capacity. Specifically, the functional decoupling between the declive of the cerebellum and the ACC seems to be compensated by increased FC of the declive with the IPL and the PCC. The exact functional role of the cerebellar declive remains to be clarified. The emerging perspective suggests that robust intrinsic mechanisms continue to drive the effort for functional adaptation in the face of neurological insult, even during the ultimate phase of AD, despite the fact that effective compensation is no longer possible.

### 4.5 Relevance to contemporary theories

Our findings provide evidence of convergence between two complementary theoretical frameworks: a) the ‘compensation-related utilization of neural circuits hypothesis’ (CRUNCH; Reuter-Lorenz and Cappell 2008) and b) the ‘network-based degeneration’ theory (Seeley et al., 2009). The integration of these two perspectives has been suggested in a recent meta-analysis of functional network alterations in AD, proposing that in the preclinical stage, compensatory mechanisms arise in the form of increased functional connectivity to counteract connectivity loss in other regions, while in more advanced stages this compensatory mechanism breaks down as a result of increasing local and global neurodegenerative pathology (Jacobs et al., 2013). We provide voxel-wise maps that depict in unprecedented detail the progressive processes that have been proposed by these contemporary theories, while highlighting their relevance to cognitive functions. NIfTI images of our findings will be shared upon request, along with the scripted analysis pipeline, to facilitate replicable Open Science practices.

### 4.6 Methodological considerations

To our knowledge, this is the first study using the p-tau/Aβ42 ratio as a regressor in a functional neuroimaging study and the first study investigating linear changes in eigenvector centrality across the pathophysiological continuum of AD. Further, it is the first time that voxel-wise eigenvector centrality has been computed in AD patients with a field of view and connectivity matrix including the cerebellum. Moreover, this is the first study investigating resting-state EC in AD voxel-wise and at 3 Tesla, thereby potentially featuring increased sensitivity in subcortical areas (Skouras et al., 2014). Moreover, previous related studies normalized datasets via basic methods, such as affine spatial transformation, despite expected morphological differences due to atrophy. Standard methods of image normalization assume similar brain anatomy across subjects that is not typical in the case of neurodegenerative diseases. Specifically, most normalization algorithms operate in vector field space and thus do not preserve topological correspondence in cases of large deformations (Avants et al., 2008). SyGN is a recent method utilizing symmetric diffeomorphic optimization and maximized cross-correlation to guarantee robust quality performance and the preservation of topological correspondence (Avants et al., 2008), proven to be top-ranking among currently available normalization methods while producing findings generalizable to new subject populations (Avants et al., 2008; Klein et al., 2009). By using SyGN and a custom anatomical template we performed the optimal normalization procedure developed for use in samples that include both intact and atrophied brains. Lastly, our data-driven approach to FC seed derivation, enabled focusing on the progressive FC changes of the brain regions predominantly implicated throughout AD, via an unbiased and information-efficient procedure.

### 4.7 Limitations and future directions

We have adopted the most straightforward interpretation of considering decreases in EC and FC to be due to AD progression and spatially linked concurrent increases to correspond to compensatory mechanisms. Moreover, we assumed that functional decoupling between two brain regions represents functional degradation and can be compensated by concurrent coupling of either decoupled region with alternative ones. Two further implicit assumptions were that the cross-sectional p-tau/Aβ42 ratio is a valid marker of position along the AD continuum and that diagnostic categories are sequential and irreversible. Although our assumptions are largely supported the scientific literature, by the current theoretical framework (Reuter-Lorenz and Cappell 2008; Seeley et al., 2009; Barulli and Stern 2013; Jacobs et al., 2013), by the corroborated meta-analyses discussed (Jacobs et al., 2013; Martinelli et al., 2013), as well by a complementary volumetric study of the same population (Gispert et al., 2015), it remains likely that the underlying mechanisms are of higher complexity. It will be important for our findings to be replicated and confirmed in longitudinal studies using larger samples, especially with regards to the preclinical group. Nevertheless, we believe that the patterns of voxel-wise connectomics revealed here can be informative for future diagnostic procedures and contribute valuable insights towards appreciating the distributed functional capacity of the human brain.

## 5. Conclusions

Preclinical functional decoupling occurs predominantly in AD-vulnerable regions (e.g. hippocampus, cerebellar lobule VI / Crus I, visual cortex, frontal pole) and each AD stage is characterized by distinct patterns of FC. Plausible mechanisms of functional compensation across the AD continuum, appear primarily in three specific forms of changes in connectivity as: a) increased EC of thalamus, BA24, BA31, BA23 and BA32 across the entire continuum; b) increased coupling of MCC with PCu and of PCC with cerebellar declive, BA18, lingual gyri and fusiform gyri in the preclinical stage and c) increased coupling of MCC with BA32, BA8 and cerebellar uvula and of PCC with inferio-anterior insula and BA47 in MCI.

## ACKNOWLEDGEMENTS

This work has received funding from the European Union’s Horizon 2020 research and innovation programme under the Marie Sklodowska-Curie action grant agreement No 707730. This work was also supported by the AETIONOMY project (Organising Mechanistic Knowledge about Neurodegenerative Diseases for the Improvement of Drug Development and Therapy) of the EU/EFPIA Innovative Medicines Initiative grant agreement No 115568 and part of the data collection was financed by the Spanish Ministry of Economy and Competitiveness ISCIII which was co-funded by the European Regional Development Fund grant number PI14/00282 (AL). JDG holds a “Ramόn y Cajal” fellowship (RYC-2013-13054).

## Supplementary material

**Supplementary Figure S1:**
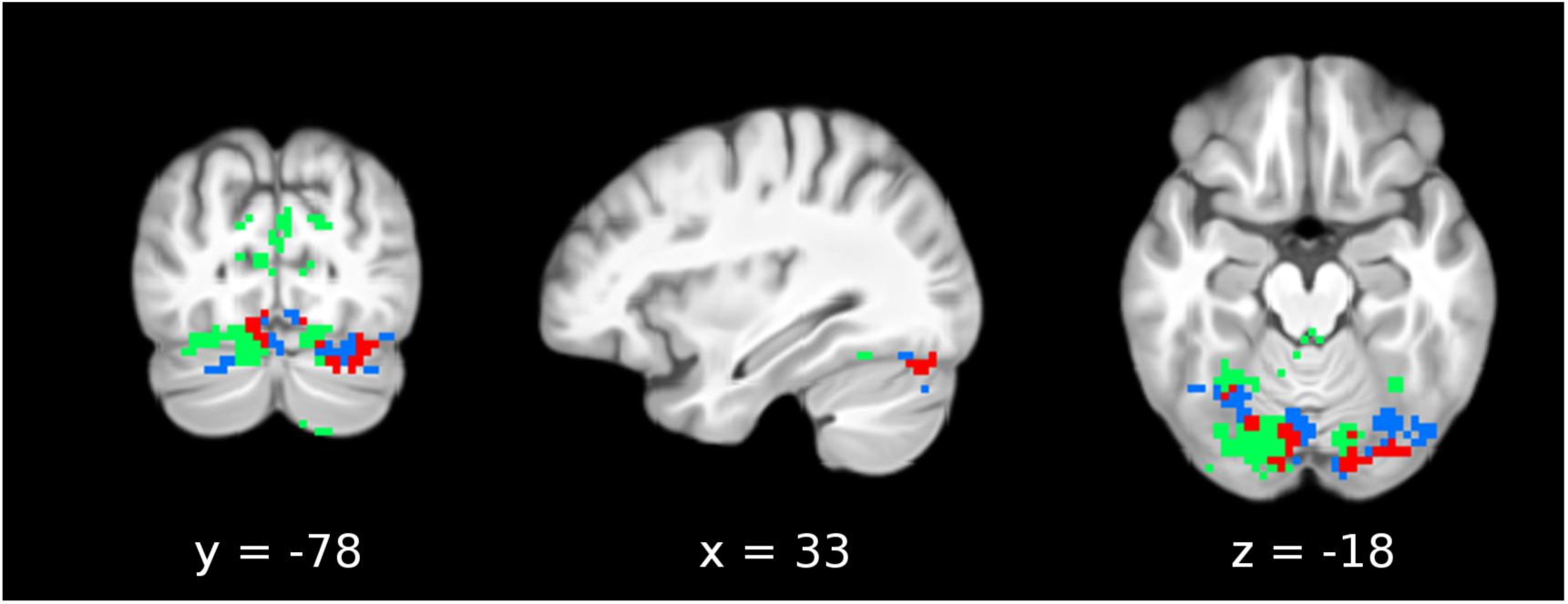
Conjunction of cerebellar FC increase with posterior cingulate and cerebellar FC decrease with IPL in the preclinical stage of Alzheimer’s disease. Blue voxels indicate significant FC decrease with IPL in the preclinical stage of AD (see Figure 3, bottom row). Red voxels indicate significant FC increase with posterior cingulate in the preclinical stage of AD (see Figure 3, second row). Green voxels indicate the overlap between red and blue clusters, suggesting their compensatory relationship. Note that many green voxels are located in the vicinity of cerebellar lobule VI and Crus I.

**Supplementary Figure S2:**
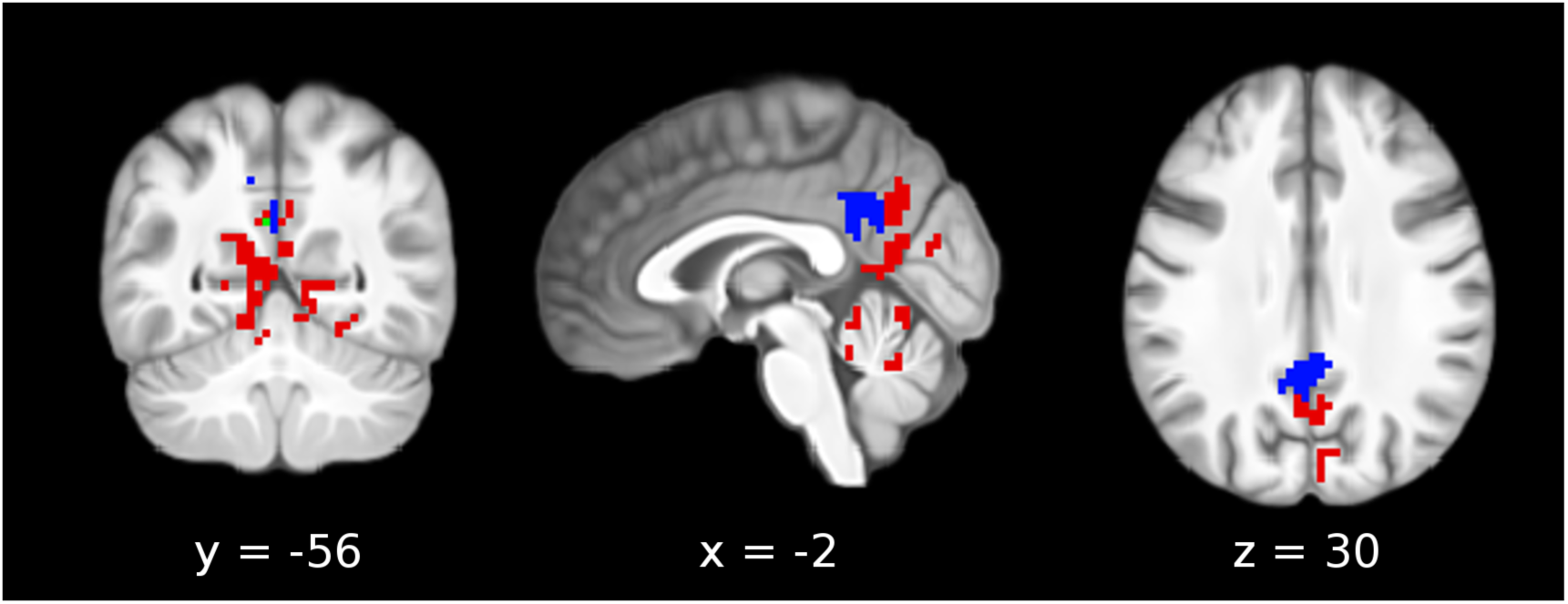
Adjacency of precuneal FC decrease with medial cingulate and precuneal FC increase with posterior cingulate in dementia due to Alzheimer’s disease. Blue voxels indicate significant FC decrease with MCC during dementia due to Alzheimer’s disease (see Figure 5, top row). Red voxels indicate significant FC increase with PCC during dementia due to AD (see Figure 5, second row). Marginal overlap was limited to 1 voxel, depicted in green.

**Supplementary table 1:** Neuropsychological tests administered.

| List of neuropsychological tests administered, organized by domain |  |
| --- | --- |
| Memory domain |  |
|  | Free and Cued Selective Reminding Test (FCSRT; Grober & Buschke, 1987) |
|  | Memory Alteration Test (M@T; Rami et al. 2007) |
| Language domain |  |
|  | Boston Naming Test (BNT; Kaplan, Goodglass, & Weintraub, 1983) |
|  | Semantic fluency (Goodglass, Kaplan, & Barresi, 2000) |
| Visual perception domain |  |
|  | Number location subtest of the VOSP battery (Warrington & James, 1991) |
| Executive functions domain |  |
|  | Trail Making Test (TMT; Reitan, 1994) |
|  | Stroop Test (Stroop, 1935) |
|  | Symbol Digit Modalities Test (SDMT; Smith, 1982) |
|  | Digit Span test of the WAIS (Wechsler, 2008) |
| Global cognition |  |
|  | Mini Mental State Evaluation (MMSE; Folstein *et al.*, 1975) |
| Premorbid intelligence |  |
|  | Spanish word accentuation test (WAT; Gomar *et al.*, 2011) |

## REFERENCES

Adriaanse SM, Wink AM, Tijms BM, Ossenkoppele R, Verfaillie SCJ, Lammertsma AA, Boellaard R, Scheltens P, van Berckel BNM, Barkhof F (2016): The Association of Glucose Metabolism and Eigenvector Centrality in Alzheimer’s Disease. Brain connectivity 6: 1–8.

Aggleton JP, Pralus A, Nelson A, Hornberger M (2016): Thalamic pathology and memory loss in early Alzheimer’s disease: moving the focus from the medial temporal lobe to Papez circuit. published online ahead of print.

Albada VS, Robinson PA (2007): Transformation of arbitrary distributions to the normal distribution with application to EEG test–retest reliability. Journal of neuroscience methods 161: 205–211.

Andrews-Hanna JR, Reidler JS, Sepulcre J, Poulin R, Buckner RL (2010): Functional-anatomic fractionation of the brain’s default network. Neuron 65: 550–562.

Avants BB, Epstein CL, Grossman M, Gee JC (2008): Symmetric diffeomorphic image registration with cross-correlation: evaluating automated labeling of elderly and neurodegenerative brain. Medical image analysis 12: 26–41.

Avants BB, Tustison N, Song G (2009): Advanced normalization tools (ANTS). Insight 2: 1–35.

Avants B, Tustison N, Song G, Cook P, Klein A, Gee J (2010): A reproducible evaluation of ANTs similarity metric performance in brain image registration. Neuroimage 54: 2033–44.

Avants B, Yushkevich P, Pluta J, Minkoff D, Korczykowski M, Detre J, et al. (2010): The optimal template effect in hippocampus studies of diseased populations. Neuroimage 49: 2457–2466.

Babiloni C, Ferretti A, Gratta DC, Carducci F (2005): Human cortical responses during one-bit delayed-response tasks: an fMRI study. Brain research bulletin 65: 383–390.

Bäckman L, Jones S, Berger AK, Laukka EJ (2005): Cognitive impairment in preclinical Alzheimer’s disease: a meta-analysis. Neuropsychology 19: 520.

Backus AR, Schoffelen JM, Szebényi S, Hanslmayr S (2016): Hippocampal-prefrontal theta oscillations support memory integration. Current Biology 26: 450–457.

Barulli D, Stern Y (2013): Efficiency, capacity, compensation, maintenance, plasticity: emerging concepts in cognitive reserve. Trends in cognitive sciences 17: 502–509.

Binnewijzend M, Adriaanse S, Flier W, Teunissen C, Munck J, Stam C, et al. (2014): Brain network alterations in Alzheimer’s disease measured by Eigenvector centrality in fMRI are related to cognition and CSF biomarkers. Human Brain Mapping 35: 2383–2393.

Binnewijzend M, Schoonheim MM (2012): Resting-state fMRI changes in Alzheimer’s disease and mild cognitive impairment. Neurobiology of aging 33: 2018–2028.

Blacker D, Albert MS, Bassett SS (1994): Reliability and validity of NINCDS-ADRDA criteria for Alzheimer’s disease: the National Institute of Mental Health Genetics Initiative. Alzheimer’s & dementia 7: 263–269.

Bonacich P (1972): Technique for analyzing overlapping memberships. Sociological methodology 4: 176–185.

Borgatti SP (2005): Centrality and network flow. Social networks 27: 55–71.

Buckner R, Sepulcre J, Talukdar T, Krienen F, Liu H, Hedden T, et al. (2009): Cortical hubs revealed by intrinsic functional connectivity: mapping, assessment of stability, and relation to Alzheimer’s disease. Journal of Neuroscience 29: 1860–73.

Cavanna AE, Trimble MR (2006): The precuneus: a review of its functional anatomy and behavioural correlates. Brain 129: 564–583.

Coupé P, Yger P, Prima S, Hellier P, Kervrann C, Barillot C (2008): An optimized blockwise nonlocal means denoising filter for 3-D magnetic resonance images. Medical Imaging, IEEE Transactions 27: 425–441.

de Souza LC, Lamari F, Belliard S, Jardel C, Houillier C, De Paz R, et al. (2011): Cerebrospinal fluid biomarkers in the differential diagnosis of Alzheimer’s disease from other cortical dementias. Journal of Neurology, Neurosurgery & Psychiatry 82: 240–6.

Dickerson BC, Bakkour A, Salat DH, Feczko E (2009): The cortical signature of Alzheimer’s disease: regionally specific cortical thinning relates to symptom severity in very mild to mild AD dementia and is detectable in in asymptomatic amyloid-positive individuals. Cerebral Cortex 19: 497–510.

Driessen M, Herrmann J, Stahl K (2000): Magnetic resonance imaging volumes of the hippocampus and the amygdala in women with borderline personality disorder and early traumatization. Arch Gen Psychiatry 57:1115–1122.

Dubois B, Hampel H, Feldman HH, Scheltens P, Aisen P, Andrieu S, Bakardjian H, Benali H, Bertram L, Blennow K, et al. (2016) Preclinical Alzheimer’s disease: definition, natural history, and diagnostic criteria. Alzheimer’s & dementia: the journal of the Alzheimer’s Association 12: 292–323.

Elman JA, Madison CM, Baker SL, Vogel JW, Marks SM, Crowley S, et al. (2016): Effects of beta-amyloid on resting state functional connectivity within and between networks reflect known patterns of regional vulnerability. Cerebral Cortex 26: 695–707.

Ethofer T, Anders S, Erb M, Herbert C, Wiethoff S (2006): Cerebral pathways in processing of affective prosody: a dynamic causal modeling study. Neuroimage 30: 580–587.

Fagan AM, Roe CM, Xiong C (2007): Cerebrospinal fluid tau/β-amyloid42 ratio as a prediction of cognitive decline in nondemented older adults. Archives of neurology 64: 343–349.

Fisher RA (1915): Frequency distribution of the values of the correlation coefficient in samples of an indefinitely large population. Biometrika: 10 (4): 507–521.

Fisher RA (1921): On the ‘probable error’ of a coefficient of correlation deduced from a small sample. Metron. 1: 3–32.

Fonov V, Evans AC, Botteron K, Almli RC, McKinstry RC, Collins LD (2011): Unbiased average age-appropriate atlases for pediatric studies. NeuroImage 54: 313–327.

Fortea J, Sala-Llonch R, Bartrés-Faz D, Lladό A (2011): Cognitively preserved subjects with transitional cerebrospinal fluid ss-amyloid 1-42 values have thicker cortex in Alzheimer’s disease vulnerable areas. Biological Psychiatry 70: 183–190.

Gallichan D, Scholz J, Bartsch A, Behrens T, Robson M, Miller K (2010): Addressing a systematic vibration artifact in diffusion-weighted MRI. Hum Brain Mapp 31: 193–202.

Ghaem O, Mellet E, Crivello F, Tzourio N, Mazoyer B (1997): Mental navigation along memorized routes activates the hippocampus, precuneus, and insula. Neuroreport 8: 739–744.

Gilbert SJ, Dumontheil I, Simons JS, Frith CD, Burgess PW (2007): Comment on “Wandering minds: the default network and stimulus-independent thought.” Science 317:43.

Gispert JD, Rami L, Sánchez-Benavides G, Falcon C, Tucholka A, Rojas S, et al. (2015): Nonlinear cerebral atrophy patterns across the Alzheimer’s disease continuum: impact of APOE4 genotype. Neurobiology of aging 36: 2687–2701.

Grober E, Buschke H (1987): Genuine memory deficits in dementia. Developmental neuropsychology 3:13–36.

Grober E, Ocepek-Welikson K, Teresi JA (2009): The free and cued selective reminding test: evidence of psychometric adequacy. Psychology Science Quarterly 51: 266–282.

Guo CC, Tan R, Hodges JR, Hu X, Sami S (2016): Network-selective vulnerability of the human cerebellum to Alzheimer’s disease and frontotemporal dementia. Brain 139: 1527–1538.

Habas C, Kamdar N, Nguyen D, Prater K (2009): Distinct cerebellar contributions to intrinsic connectivity networks. The Journal of Neuroscience 29: 8586–8594.

Hesling I, Clément S, Bordessoules M, Allard M (2005): Cerebral mechanisms of prosodic integration: evidence from connected speech. Neuroimage 24: 937–947.

Hill AC, Laird AR, Robinson JL (2014): Gender differences in working memory networks: A BrainMap meta-analysis. Biological psychology 102: 18–29.

Iglόi K, Doeller CF, Paradis A-L, Benchenane K, Berthoz A, Burgess N, et al. (2015) Interaction between hippocampus and cerebellum crus I in sequence-based but not place-based navigation. Cerebral Cortex 25: 4146–4154.

Jack C, Wiste H, Weigand S, Knopman D, Lowe V, Vemuri P, et al. (2013): Amyloid-first and neurodegeneration-first profiles characterize incident amyloid PET positivity. Neurology 81: 1732–1740.

Jack C, Knopman D, Jagust W, Petersen R, Weiner M, Aisen P, et al. (2013): Tracking pathophysiological processes in Alzheimer’s disease: an updated hypothetical model of dynamic biomarkers. Lancet Neurology 12: 207–16.

Jacobs H, Radua J, Lückmann HC (2013): Meta-analysis of functional network alterations in Alzheimer’s disease: toward a network biomarker. Neuroscience & Biobehavioral Reviews 37: 753–65.

Janata P (2009): The neural architecture of music-evoked autobiographical memories. Cerebral Cortex 19: 2579–2594.

Kapur S, Craik FI, Tulving E (1994): Neuroanatomical correlates of encoding in episodic memory: levels of processing effect. PNAS 91: 2008–2011.

Klein A, Andersson J, Ardekani BA, Ashburner J, Avants B, Chiang M-CC, et al. (2009): Evaluation of 14 nonlinear deformation algorithms applied to human brain MRI registration. Neuroimage 46: 786–802.

Koelsch S, Skouras S (2014): Functional centrality of amygdala, striatum and hypothalamus in a “small-world” network underlying joy: An fMRI study with music. Human brain mapping 35: 3485–3498.

Krienen FM, Buckner RL (2009): Segregated fronto-cerebellar circuits revealed by intrinsic functional connectivity. Cerebral cortex 19: 2485–97.

Kurth F, Zilles K, Fox PT, Laird AR (2010): A link between the systems: functional differentiation and integration within the human insula revealed by meta-analysis. Brain Structure and Function 214: 519–534.

Leal SL, Landau SM, Bell RK, Jagust WJ (2017): Hippocampal activation is associated with longitudinal amyloid accumulation and cognitive decline. eLife 6: e22978.

Lemieux L, Salek-Haddadi A, Lund TE, Laufs H (2007): Modelling large motion events in fMRI studies of patients with epilepsy. Magnetic resonance imaging 25:894–901.

Li G, Sokal I, Quinn JF, Leverenz JB, Brodey M, Schellenberg GD, Kaye JA, Raskind MA, Zhang J, Peskind ER, et al. (2007) CSF tau/Aβ42 ratio for increased risk of mild cognitive impairment A follow-up study. Neurology 69: 631–639}.

Lohmann G, Müller K, Bosch V, Mentzel H, Hessler S (2000): LIPSIA-Leipzig Image Processing and Statistical Inference Algorithms.

Lohmann G, Neumann J, Müller K (2008): The multiple comparison problem in fmri - A new method based on anatomical priors. Proceedings of the First Workshop on Analysis of Functional Medical Images.

Lohmann G, Margulies DS, Horstmann A, Pleger B, Lepsien J, Goldhahn D, et al. (2010): Eigenvector centrality mapping for analyzing connectivity patterns in fMRI data of the human brain. PloS one 5: e10232.

Maddalena A, Papassotiropoulos A, Müller-Tillmanns B, Jung HH, Hegi T, Nitsch RM, Hock C (2003): Biochemical diagnosis of Alzheimer disease by measuring the cerebrospinal fluid ratio of phosphorylated tau protein to β-amyloid peptide42. Archives of neurology 60: 1202–1206.

Martinelli P, Sperduti M, Piolino P (2013): Neural substrates of the self-memory system: New insights from a meta - analysis. Human Brain Mapping 34: 1515–1529.

McDowell JE, Dyckman KA, Austin BP, Clementz BA (2008): Neurophysiology and neuroanatomy of reflexive and volitional saccades: evidence from studies of humans. Brain and cognition 68: 255–270.

Molinuevo J, Gispert J, Dubois B, Heneka M, Lleo A, Engelborghs S, et al. (2013): The AD-CSF-index discriminates Alzheimer’s disease patients from healthy controls: a validation study. Journal of Alzheimer’s Disease 36: 67–77.

Molitor RJ, Ko PC, Ally BA (2015): Eye movements in Alzheimer’s disease. Journal of Alzheimer’s Disease 44: 1–12.

Menon V, Uddin LQ (2010): Saliency, switching, attention and control: a network model of insula function. Brain Structure and Function 214:655–67.

Palmqvist S, Schöll M, Strandberg O, Mattsson N, Stomrud E, Zetterberg H, Blennow K, Landau S, Jagust W, Hansson O (2017): Earliest accumulation of β-amyloid occurs within the default-mode network and concurrently affects brain connectivity. Nature Communications 8: 1214.

Pihlajamäki M, Sperling RA (2008): fMRI: use in early Alzheimer’s disease and in clinical trials. Future Neurology 3: 409–421.

Rami L, Molinuevo JL, Sanchez - Valle R (2007): Screening for amnestic mild cognitive impairment and early Alzheimer’s disease with M@T (Memory Alteration Test) in the primary care population. International journal of geriatric psychiatry 22: 294–304.

Ranganath C, Johnson MK, D’Esposito M (2003): Prefrontal activity associated with working memory and episodic long-term memory. Neuropsychologia 41: 378–389.

Raichle ME, MacLeod AM, Snyder AZ, Powers WJ, Gusnard D, et al (2001): A default mode of brain function. PNAS 98: 676–82.

Gusnard DA, Raichle ME (2001): Searching for a baseline: functional imaging and the resting human brain. Nature Reviews Neuroscience 2: 685–694.

Reuter-Lorenz, P.A. and Cappell, K.A., 2008. Neurocognitive aging and the compensation hypothesis. Current directions in psychological science, 17(3), pp. 177–182.

Rowe CC, Ellis KA, Rimajova M, Bourgeat P, Pike KE, Jones G, Fripp J, Tochon-Danguy H, Morandeau L, O’Keefe G, Price R (2010): Amyloid imaging results from the Australian Imaging, Biomarkers and Lifestyle (AIBL) study of aging. Neurobiology of aging 31:1275–83.

Sadananthan, S.A., Zheng, W., Chee, M.W. and Zagorodnov, V., 2010. Skull stripping using graph cuts. NeuroImage, 49(1), pp.225–239.

Schnell K, Dietrich T, Schnitker R, Daumann J (2007): Processing of autobiographical memory retrieval cues in borderline personality disorder. Journal of affective disorders 97: 253–259.

Schoonheim MM, Geurts JJG, Wiebenga OT, De Munck JC, Polman CH, Stam CJ, Barkhof F, Wink AM (2014): Changes in functional network centrality underlie cognitive dysfunction and physical disability in multiple sclerosis. Multiple Sclerosis Journal 20: 1058–1065.

Seeley WW, Crawford RK, Zhou J, Miller BL, Greicius MD (2009): Neurodegenerative diseases target large-scale human brain networks. Neuron 62: 42–52.

Sheline Y, Raichle M (2013): Resting State Functional Connectivity in Preclinical Alzheimer’s Disease. Biological Psychiatry 74: 340–347.

Shulman GL, Fiez JA, Corbetta M, Buckner RL, Miezen FM, Raichle ME et al (1997): Common blood flow changes across visual tasks, II: decreases in cerebral cortex. J Cogn Neurosci 9: 648–63.

Skouras S, Gray M, Critchley H, Koelsch S (2014): Superficial Amygdala and Hippocampal Activity During Affective Music Listening at 3 T but not 1.5 T fMRI. NeuroImage 101: 364–369.

Skouras S, Gispert JD, Molinuevo JL (2018): The Crus exhibits stronger functional connectivity with executive network nodes than with the default mode network. Brain 141: e24.

Söderlund H, Moscovitch M, Kumar N, Mandic M (2012): As time goes by: Hippocampal connectivity changes with remoteness of autobiographical memory retrieval. Hippocampus 22: 670–679.

Sperling RA, Aisen PS, Beckett LA, Bennett DA, Craft S, Fagan AM, Iwatsubo T, Jack CR, Kaye J, Montine TJ, Park DC (2011): Toward defining the preclinical stages of Alzheimer’s disease: Recommendations from the National Institute on Aging-Alzheimer’s Association workgroups on diagnostic guidelines for Alzheimer’s disease. Alzheimer’s & dementia 7: 280–292.

Spreng RN, Stevens WD, Chamberlain JP, Gilmore AW, Schacter DL (2010). Default network activity, coupled with the frontoparietal control network, supports goal-directed cognition. Neuroimage, 53(1), pp.303–317.

Szirmai I, Vastagh I, Szombathelyi É, Kamondi A (2002): Strategic infarcts of the thalamus in vascular dementia. Journal of the neurological sciences 203: 91–97.

Teipel S, Drzezga A, Grothe MJ, Barthel H (2015): Multimodal imaging in Alzheimer’s disease: validity and usefulness for early detection. The Lancet Neurology 14:1037–53.

Tustison NJ, Avants BB, Cook PA (2013): The ANTs cortical thickness processing pipeline. SPIE Medical Imaging. International Society for Optics and Photonics.

Tustison NJ, Avants BB (2013): Explicit B-spline regularization in diffeomorphic image registration. Frontiers in neuroinformatics 7: 39.

Tustison N, Avants B, Cook P, Zheng Y, Egan A, Yushkevich P, et al. (2010): N4ITK: improved N3 bias correction. Ieee T Med Imaging 29:1310–20.

Volz KG, Schubotz RI, von Cramon DY (2004): Why am I unsure? Internal and external attributions of uncertainty dissociated by fMRI. Neuroimage 21: 848–857.

Wager TD, Smith EE (2003): Neuroimaging studies of working memory. Cognitive, Affective, & Behavioral Neuroscience 3: 255–274.

Wang L, Zang Y, He Y, Liang M, Zhang X, Tian L, et al. (2006): Changes in hippocampal connectivity in the early stages of Alzheimer’s disease: Evidence from resting state fMRI. Neuroimage 31: 496–504.

Wesson J, Luchins J (1992): An empirical evaluation of the Global Deterioration Scale for staging Alzheimer’s disease. Am. J. Psychiatry 149:190–194.

Zhang JX, Leung HC, Johnson MK (2003): Frontal activations associated with accessing and evaluating information in working memory: an fMRI study. Neuroimage 20: 1531–1539.

Zhou B, Liu Y, Zhang Z, An N, Yao H, Wang P, et al. (2013): Impaired functional connectivity of the thalamus in Alzheimer’s disease and mild cognitive impairment: a resting-state fMRI study. Current Alzheimer Research 10: 754–766.

## References for supplementary table 1

Fosltein MF, Folstein SE, McHugh PR. Mini-mental state. A practical method for grading the cognitive state of patients for the clinician. J Psychiatr Res. 1975;12:189–98.

Gomar JJ, Ortiz-Gil J, McKenna PJ, Salvador R, Sans-Sansa B, Sarrό S, Guerrero A, Pomarol-Clotet E. Validation of the Word Accentuation Test (TAP) as a means of estimating premorbid IQ in Spanish speakers. Schizophrenia research. 2011;128:175–6.

Goodglass H, Kaplan E, Barresi B. Boston Diagnostic Aphasia Examination-(BDAE-3) San Antonio. TX: Psychological Corporation. 2000.

Grober E, Buschke H. Genuine memory deficits in dementia. Developmental neuropsychology 1987; 3:13–36.

Kaplan E, Goodglass H, Weintraub S. Boston Naming Test (BNT). Manual (2nd edition). Philadelphia: Lea and Fabiger. 1983.

Rami L, Molinuevo JL, Sanchez-Valle R. Screening for amnestic mild cognitive impairment and early Alzheimer’s disease with M@T (Memory Alteration Test) in the primary care population. International journal of geriatric psychiatry 2007. 22: 294–304.

Reitan RM, Wolfson D. The Halstead–Reitan neuropsychological test battery for Adults—Theoretical, methodological, and validational bases. Neuropsychological assessment of neuropsychiatric and neuromedical disorders. 2009;1.

Smith A. Symbol digit modalities test (SDMT) manual (revised) Western Psychological Services. Los Angeles. 1982. Stroop JR. Studies of interference in serial verbal reactions. Journal of experimental psychology. 1935;18:643.

Warrington EK, James M. VOSP Visual Object and Space Perception Test Battery. Bury St Edmunds: TVTC Thames Valley Test Company. 1991.

Wechsler D. Wechsler Adult Intelligence Scale–Fourth Edition (WAIS-IV). 2014.

